# Global brain activity and its coupling with cerebrospinal fluid flow is related to tau pathology

**DOI:** 10.1101/2023.09.12.557492

**Authors:** Feng Han, JiaQie Lee, Xi Chen, Jacob Ziontz, Tyler Ward, Susan M Landau, Suzanne L Baker, Theresa M Harrison, William J Jagust, the Alzheimer’s Disease Neuroimaging Initiative

## Abstract

Amyloid-β (Aβ) and tau deposition constitute Alzheimer’s disease (AD) neuropathology. Cortical tau deposits first in the entorhinal cortex and hippocampus and then propagates to neocortex in an Aβ-dependent manner. Tau also tends to accumulate earlier in higher-order association cortex than in lower-order primary sensory-motor cortex. While previous research has examined the production and spread of tau, little attention has been paid to its clearance. Low-frequency (<0.1 Hz) global brain activity during the resting state is coupled with cerebrospinal fluid (CSF) flow and potentially reflects glymphatic clearance. Here we report that tau deposition in subjects with evaluated Aβ, accompanied by cortical thinning and cognitive decline, is strongly associated with decreased coupling between CSF flow and global brain activity. Substantial modulation of global brain activity is also manifested as propagating waves of brain activation between higher- and lower-order regions, resembling tau spreading. Together, the findings suggest an important role of resting-state global brain activity in AD tau pathology.

**One Sentence Summary:** Resting-state global brain activity affects tau deposition through the potential involvement of a glymphatic clearance function.

## Introduction

Alzheimer’s disease (AD) is pathologically characterized by the extracellular accumulation of amyloid-β (Aβ) plaques and the intracellular accumulation of hyperphosphorylated tau in the form of neurofibrillary tangles (*1–4*). Converging evidence has shown that Aβ accelerates tau phosphorylation and promotes tau aggregation and oligomerization, while Aβ toxicity is dependent on tau (*5–7*). Tau plays an especially critical role in cognitive decline (*8*), brain atrophy, particularly cortical thinning (*9, 10*) and neuronal and synaptic loss (*8, 11*). For example, a recent study has shown that tau burden measured with positron emission tomography (PET) predicts the severity and topography of subsequent cortical atrophy in AD patients (*12*).Recently, tau has received special attention as a potential therapeutic target of AD. The deposition of tau aggregates in AD follows a stereotypical pattern, beginning in the entorhinal cortex and hippocampus and then propagating to distinct regions of interest (ROIs) that have been characterized in postmortem and imaging studies as Braak Stages (*13, 14*). Recent studies have characterized tau deposition as occurring later in lower-order primary sensory-motor (SM) regions but earlier in higher-order association regions that can be captured through the examination of network, such as the default mode network (DMN) and frontoparietal network (FPN) (*15, 16*). The neural mechanism underlying such a stereotyped pattern of tau accumulation remains elusive but is likely multifactorial. Current hypotheses have attributed tau spreading to neural activity (*17, 18*), and a “prion-like” mechanism manifested as abnormal tau seeds transferring through anatomically (*19–23*) and functionally (*24–26*) connected brain regions.

Beyond the accumulation of tau pathology, its clearance may be anti-correlated with deposition and thus has received increasing attention (*27, 28*). For example, recent studies have identified the critical role of glymphatic function in clearing brain wastes, including Aβ and tau, via a pathway involving cerebrospinal fluid (CSF) flow, and the exchange between CSF and interstitial fluid (ISF), as well as ISF solutes (*27, 29, 30*). In mice, the sleep-wake cycle has been found to regulate ISF tau, and sleep deprivation can significantly increase ISF and CSF tau as well as tau spreading (*31*), presumably due to inadequate sleep-dependent glymphatic clearance (*29*). This sleep-dependent feature links glymphatic function with spontaneous low-frequency (< 0.1 Hz) resting-state global brain activity assessed with the global blood-oxygen-level-dependent (global BOLD, gBOLD) signal in functional MRI that increases during light sleep or low arousal states (*32–36*). More importantly, CSF movement, a key determinant of glymphatic function, is coupled with gBOLD signal during both sleep (*37*) and wakefulness (*38*). The coupling strength between CSF inflow and gBOLD fMRI signal has recently been proposed as a method to evaluate glymphatic function, and found to be related to cortical Aβ in an early AD cohort (*38, 39*), cognitive decline in AD and Parkinson’s disease patients (*38, 40*), and aging (*41*).

Of note, a recent study has linked the early accumulation of cortical Aβ in higher-order association regions to the local brain activity there and its coupling to CSF inflow (*39*), which also suggests the potential role of both global and regional low-frequency (< 0.1 Hz) neural activity in driving glymphatic clearance of Aβ. Consistent with this notion, the spread of Aβ and tau aggregates, showing similar although not identical predominance for higher-order cognitive networks over primary sensory-motor networks (*15, 16, 42*), has been hypothesized to follow the direction of glymphatic inflow (*43*). Resting-state global brain activity, measured by gBOLD or brain-wide electrophysiological signals, is strongly associated with lower-order sensory-motor activity (*44–46*), resembling the opposite spatial pattern of tau spreading (*42*). In addition, global brain activity often takes the form of dynamic propagating waves (around gBOLD peaks) between higher-order association and lower-order sensory-motor regions (*39, 47, 48*), also showing spatial correspondence with tau spreading. These dynamic propagating waves at gBOLD peaks have also been found to attenuate with the decrease of Aβ42 in CSF in the earliest AD stage (*39*), which further suggests the close link between AD pathology and the gBOLD-related propagating waves, presumably through glymphatic function. Together, these findings point to key questions: Does coupling between the global brain activity and CSF movement (gBOLD-CSF coupling), assessing glymphatic function, explain inter-subject variability of tau? Are propagating waves around gBOLD peaks related to glymphatic function, and do they explain the spatial pattern of tau deposition and spreading?

To address these hypothetical questions, we examined multimodal data from the Alzheimer’s Disease Neuroimaging Initiative-3 (ADNI3) to investigate the relationship among the coupling between gBOLD and CSF inflow fMRI signals, tau distribution measured with PET, cortical thickness, cognitive function, and dynamic propagating waves.

## Results

### Cohort demographics

We analyzed resting-state fMRI (rsfMRI), structural MRI, and PET data from 115 participants (72.5 ± 7.8 years; 60 females) in the ADNI-3 project (*49, 50*). The participants included 6 AD patients, 42 with mild cognitive impairment (MCI), 5 subjects with subjective memory concern (SMC), and 62 healthy controls. These subjects were selected based on the availability of tau-PET, Aβ-PET, rsfMRI (advanced sequence only; repetition time [TR] = 0.607 sec or s), and cortical thickness. Cortical [^18^F]florbetaben (FBB) and [^18^F]florbetapir (FBP) standardized uptake value ratios (SUVRs) were used to separate the Aβ+ (FBB > 1.08 SUVR; FBP > 1.11 SUVR) and Aβ- subjects following previous studies (*51, 52*). The diagnostic information (cognitively unimpaired: SMC and control; cognitively impaired: AD and MCI) was also employed to further define the sub-groups reflecting AD progression, including unimpaired Aβ-, unimpaired Aβ+, and impaired Aβ+ (see **Table 1** for details). In addition, 20 impaired Aβ- subjects were included to create comparable samples of Aβ- and Aβ+ individuals for fMRI-based glymphatic comparison measures, but the impaired Aβ- individuals were not included in our main analyses of glymphatic function-tau associations). In general, Aβ+ and Aβ- subjects were different (*p* < 0.001) in age and cognitive function, i.e., Montreal Cognitive Assessment (MoCA), but with a similar sex ratio between males and females. The unimpaired Aβ- subjects were also relatively younger and had higher MoCA scores compared with the impaired Aβ+ subjects (both *p* < 3.7×10^-3^).

**Table 1.** Participant characteristics. Data represent the “mean (standard deviation)” unless otherwise indicated. Two-sample t-test was applied to compare the continuous measures, while a Fisher exact test between groups was used for the sex ratio. Aβ+: cortical AV45–Aβ > 1.11 SUVR or cortical FBB–Aβ > 1.08 SUVR; SUVR: standardized uptake value ratio referring to whole cerebellum reference region; M/F: male/female; AD, Alzheimer’s disease group; MCI: mild cognitive impairment; SMC: subjective memory concern; Unimpaired diagnosis group: SMC and Control subjects; Impaired group: AD and MCI subjects.

| **N=115** | **Aβ- (N = 66)** | **Aβ+ (N = 49)** |  | **P-value** |  |  |
| --- | --- | --- | --- | --- | --- | --- |
| Age | 70.0 (7.6) | 75.8 (6.9) |  | **$5.5 \times 10^{-5}$** |  |  |
| Sex (M/F) | 32/34 | 23/26 |  | 1.0 |  |  |
| Diagnosis<br>(AD:MCI:SMC:<br>Control) | 0:20:1:45 | 6:22:4:17 |  | - |  |  |
| MoCA | 25.3 (2.9)<br>(3 N/A) | 22.3 (4.8)<br>(1 N/A) |  | **$1.0 \times 10^{-4}$** |  |  |
|  | **Stage1 (S1):**<br>**Unimpaired Aβ-**<br>**(N = 46)** | **Stage2 (S2):**<br>**Unimpaired Aβ+**<br>**(N = 21)** | **Stage3 (S3):**<br>**Impaired Aβ+**<br>**(N = 28)** | **P-value** |  |  |
|  |  |  |  | **S2 vs S1** | **S3 vs S1** | **S3 vs S2** |
| Age | 69.4 (7.8) | 76.7 (5.5) | 75.1 (7.9) | **$2.9 \times 10^{-4}$** | **$3.7 \times 10^{-3}$** | 0.42 |
| Sex (M/F) | 17/29 | 6/15 | 17/11 | 0.59 | 0.057 | **0.042** |
| Diagnosis | 0:0:1:45 | 0:0:4:17 | 6:22:1:45 | - | - | - |
| MoCA | 25.1 (2.8)<br>(2 N/A) | 25.5 (2.7) | 19.8 (4.7)<br>(1 N/A) | 0.60 | **$7.2 \times 10^{-8}$** | **$8.0 \times 10^{-6}$** |

### fMRI-based glymphatic measure is closely related to tau across A***β***+ subjects

CSF inflow rsfMRI signal was negatively correlated with gBOLD signal with a +4.856 sec time-lag (**Fig. S1C**; with a representative example in **Fig. S1, A** and **B**), which confirmed the strong coupling between the two suggested by previous work (*38*). The gBOLD-CSF coupling relevant to glymphatic function (*38*) was positively correlated with tau in most cortical regions across all subjects, i.e., subjects with more cortical tau deposition had weaker (less negative) coupling (**Fig. 1A**), similar to the trend noted previously of reduced coupling strength in subjects with more Aβ (*38*). This association was significant for regional tau deposition in the Braak V-VI ROI (isocortical regions), Braak III-IV ROI (limbic area; see ref. (*14*) for detailed cortical regions), and temporal meta-ROI (*53*) across the whole cohort (all *r* > 0.21, all *p* < 0.026; N = 115; **Fig. 1B**). Results were similar in the Braak I ROI/entorhinal cortex although not significant (*r* = 0.14, *p =* 0.12). The strong coupling-tau relationship was evident for Aβ+ and impaired Aβ+ subjects (**Fig. 1, C-F**), but not for subjects who were Aβ- or unimpaired Aβ+. (**Fig. S2**). Of note, tau remained strongly correlated with the gBOLD-CSF coupling after adjusting for the subjects’ motion level (**Fig. S3**).

**Fig. 1.**
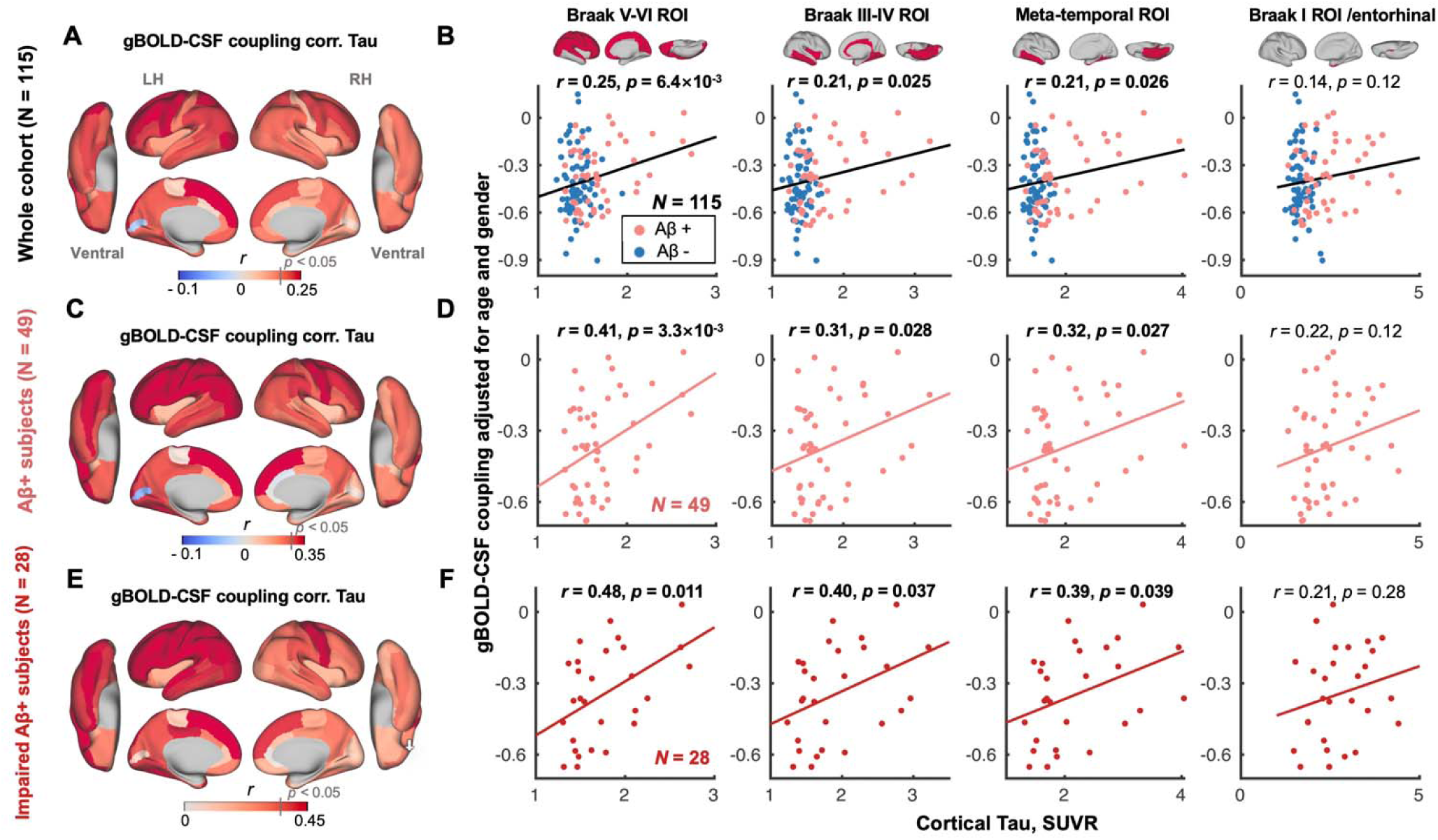
gBOLD-CSF coupling is correlated with the cortical tau across the whole cohort and Aβ+ subjects. (**A**) gBOLD-CSF coupling was positively correlated with tau in most cortical regions across all subjects, i.e., subjects with weaker (less negative) coupling had more tau deposition. (**B**) gBOLD-CSF coupling (strength) significantly decreased with more tau deposition in Braak V-VI, Braak III-IV, and temporal meta-ROI across the whole cohort (all *r* > 0.21, all *p* < 0.026; N = 115). This association was similar, although not significant (*r* = 0.14, *p* = 0.12), for tau in Braak I (entorhinal cortex). (**C-F**) These associations between gBOLD-CSF coupling and regional tau were also evident among Aβ+ and/or specifically impaired Aβ+ subjects (all *r* > 0.31, all *p* < 0.039; in Braak V-VI, Braak III-IV, and temporal meta-ROI), but not for the rest of subjects (**Fig. S2**). Aβ+: cortical AV45–Aβ > 1.11 SUVR or cortical FBB–Aβ > 1.08 SUVR. Impaired group: AD and MCI subjects. Each point represents one subject. The linear regression line was estimated based on the linear least-squares fitting (the same hereinafter unless noted otherwise).

### Tau partially mediates the strong association between gBOLD-CSF coupling and cortical thickness

We next examined the coupling-thickness association, as tau is closely associated with cortical atrophy and ultimately leads to cognitive decline (*12, 54*). Similar to the coupling-tau links above, gBOLD-CSF coupling was also significantly related to cortical thickness in widespread cortical regions, including the Braak V-VI ROI, Braak III-IV ROI, and temporal meta-ROI, across the entire group of subjects, and more specifically among the impaired Aβ+ subjects (**Fig. 2**), rather than other groups (**Fig. S4**).

**Fig. 2.**
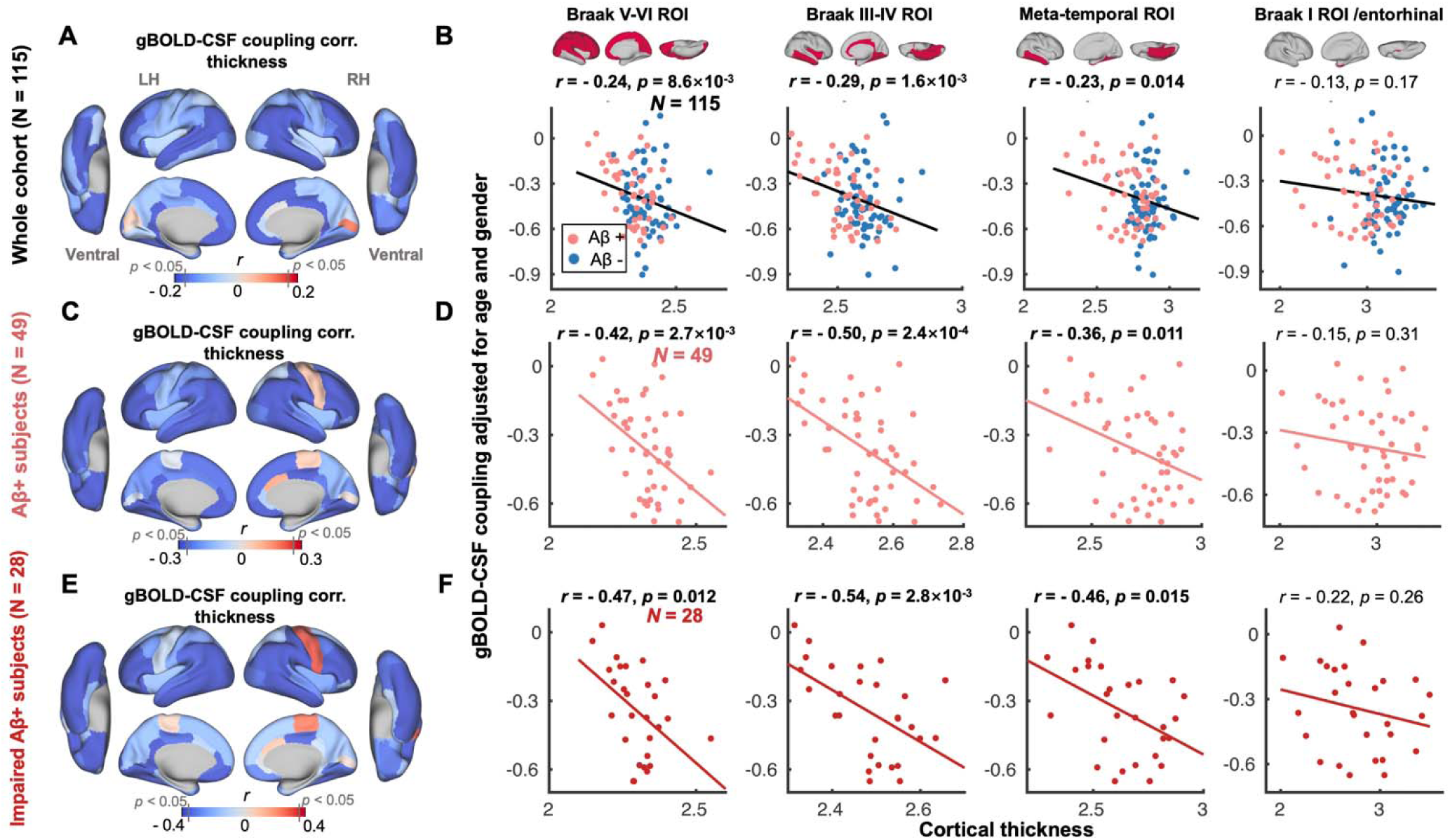
gBOLD-CSF coupling is correlated with cortical thickness across the whole cohort and Aβ+ subjects. (**A**) Subjects with weaker (less negative) gBOLD-CSF coupling had thinner cortices in the majority of brain regions, especially in the frontal, parietal, and temporal lobes, including default mode network (DMN) and fronto-parietal network (FPN). (**B**) The gBOLD-CSF coupling strength significantly decreased with thinner cortices in Braak V-VI, Braak III-IV, and temporal meta-ROIs across the whole cohort (all *r* < - 0.23, all *p* < 0.014; N = 115). A similar but not significant coupling-thickness association was found in the entorhinal region (*r* = - 0.13, *p* = 0.17). (**C-F**) Among Aβ+ subjects, particularly impaired Aβ+ ones, the coupling-thickness remained striking (all *r* < - 0.36, all *p* < 0.015 in Braak V-VI, Braak III-IV, and temporal meta-ROI in **D** and **F**) while this was not the case in other participants (**Fig. S4**). Each point represents one subject.

Cortical tau and thickness show a similar spatial distribution pattern averaged across the entire cohort and when contrasting Aβ+ and Aβ- subjects (both *p <* 1.8×10^-4^; **Fig. 3, A** and **B**). Cortical tau and thickness were also closely associated across subjects (all *p <* 0.05; **Fig. S5**). Given the corresponding spatial pattern as well as the predictive role of tau in brain atrophy (*12*), we used mediation analysis to examine whether tau in Braak V-VI ROI, Braak III-IV ROI, and the temporal meta-ROI mediated the observed relationship between gBOLD-CSF coupling and cortical thickness in the same regions. We found that tau in the three ROIs played a significant role in mediating the coupling-thickness link across the whole cohort and the Aβ+ group (*p <* 0.05; **Fig. 3, C** and **D**), although the tau in Braak V-VI ROI was a marginally significant mediator among Aβ+ subjects (*p=* 0.09). The mediation effect was less robust in impaired Aβ+ subjects (**Fig. 3E**; only significant in the meta-temporal region, *p=* 0.02), which may be attributed to the limited sample size of this group (N = 28). We also examined whether coupling mediates the tau-thickness relationship. However, the effect size of this mediation was smaller and only significant for limited ROIs among all Aβ+ and impaired Aβ+ subjects (**Fig. S6**). Together, these results suggest that tau is a mediator of the association between glymphatic function, reflected by gBOLD-CSF coupling, and cortical thickness.

**Fig. 3.**
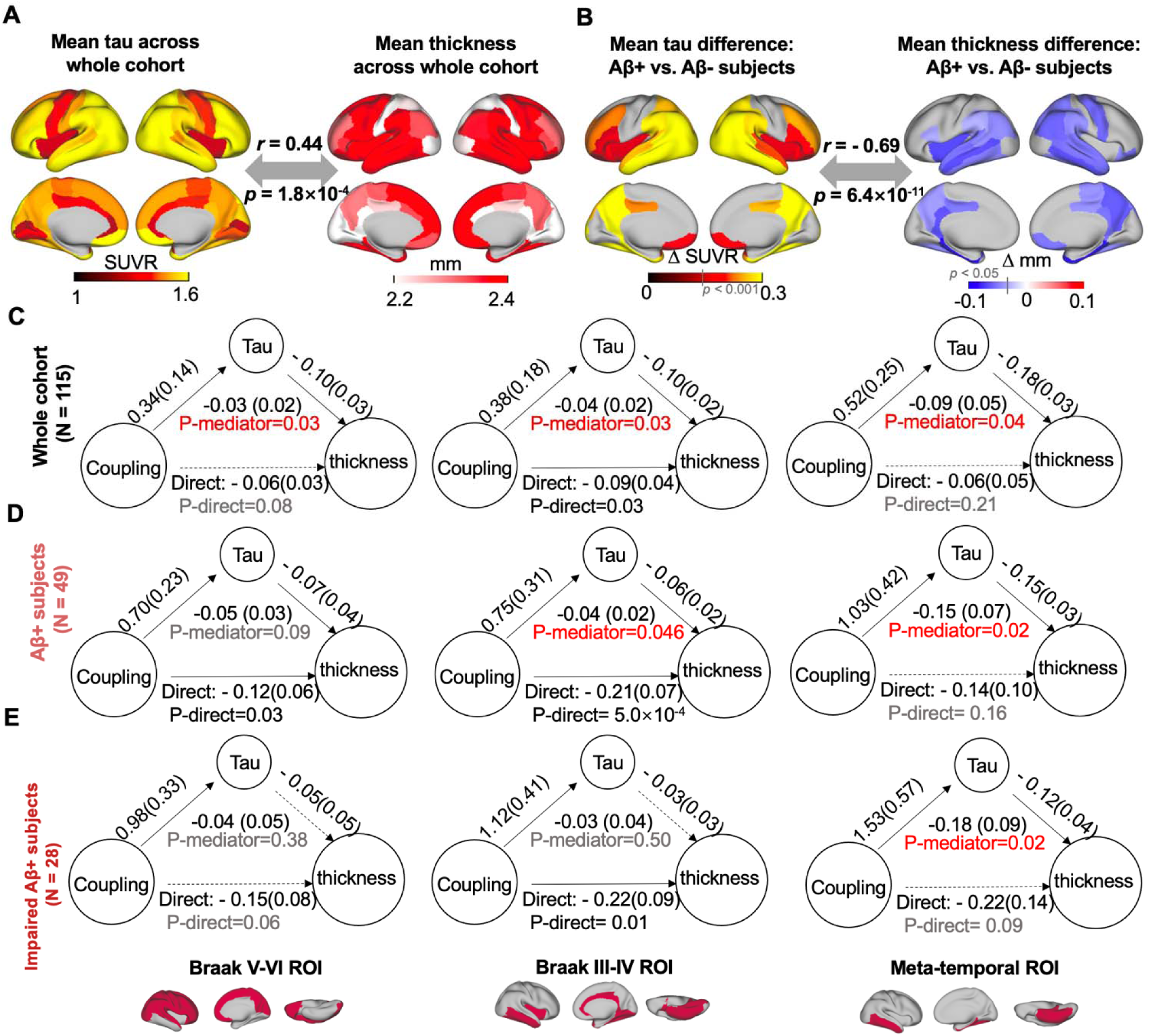
The association between coupling metrics and thickness is mediated by tau. (**A-B**) The averaged map of cortical tau and thickness across all subjects (*r* = 0.44, *p* = 1.8×10^-4^), as well as the difference between Aβ- and Aβ+ subjects (*r* = - 0.69, *p* = 6.4×10^-11^), indicative of tau and atrophy distributions were similar. This close association between tau pathology and atrophy is also consistent with their significant correlation across subjects (**Fig. S5**). (**C-E**) Across the entire cohort and Aβ+ subjects, tau mediated the significant association in Fig. 2 between gBOLD-CSF coupling and thickness throughout cortex, including Braak V-VI, Braak III-IV, and meta-temporal area (*p* < 0.046, except for the marginally significant relationship [*p* = 0.090] in Braak V-VI among the Aβ+ subjects), whereas the meditation effect was weaker for impaired Aβ+ ones, which might be attributed to the limited sample size.

Given the close association between cortical thickness and MoCA scores (**Fig. S7**), we also found that tau partially mediated the association between the coupling metrics and MoCA (**Fig. S8**), indicating a role of glymphatic clearance of tau in cognitive decline.

### Preferential tau deposition in high-order regions is related to the gBOLD-CSF coupling decrease

In the pathophysiology of AD, tau aggregates earlier in high-order association regions than in lower-order sensory ones (*15, 16, 55, 56*). We next examined how this preferential tau deposition in high-order regions is related to gBOLD-CSF coupling. Previous studies have examined patterns of functional connectivity covariance and depicted continuous and slow brain hierarchy changes across cortices with the concept of “principal gradient” (PG; **Fig. 4A**) anchored at one end by lower-order primary sensory/motor regions and at the other end by higher-order association areas (*47, 57*). Consistent with previous studies (*15, 16, 55, 56*), we demonstrated preferential tau deposition in higher-order regions by spatially correlating the PG pattern with the group-difference in tau between Aβ+ and Aβ- subjects or group-mean tau maps across Desikan-Killiany-Tourville (DKT-68) parcels (*58*). Both significant associations (both *r >* 0.26, *p <* 0.032; **Fig. 4B**) suggested that tau deposition and progression increases with the hierarchy across the entire cortex.

**Fig. 4.**
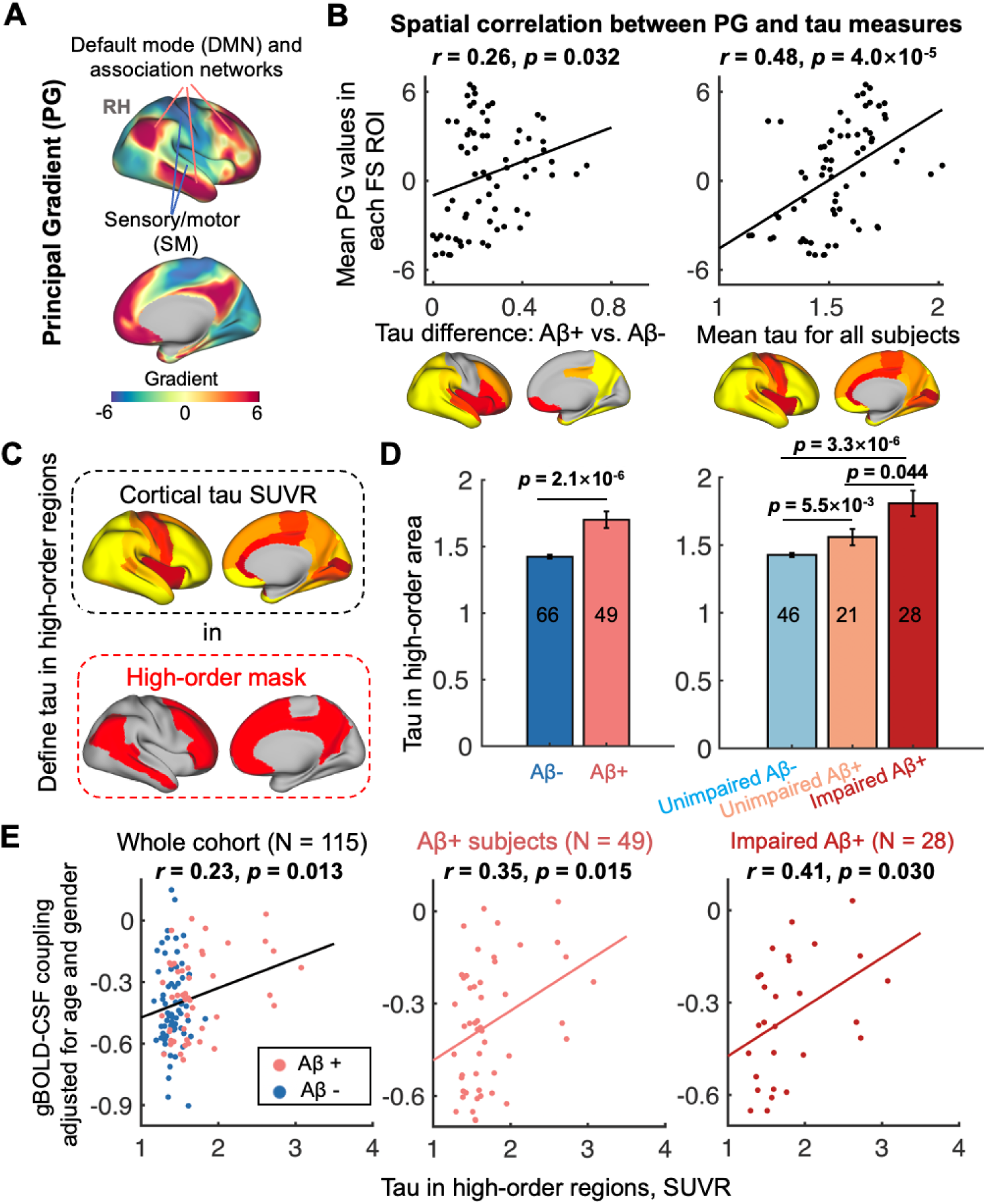
Preferential tau deposition in higher-order regions is related to coupling decrease. (**A**) A principal gradient (PG) of functional connectivity identified by a previous study (*57*) that classified cortical hierarchies by examining similarity of connectivity was employed to quantify regional cortical hierarchy. (**B**) Cortical regions (DKT-parcels) with higher tau burden, comparing Aβ- and Aβ+ individuals, were correlated with higher cortical hierarchy (*r* = 0.26, *p* = 0.032). Similarly, among all subjects, higher order regions had more tau deposition (*r* = 0.48, *p* = 4.0×10^-5^). (**C**) Two association networks with the highest PG scores, i.e., DMN and FPN (*111*), were combined and served as a higher-order mask (*39*) and used to extract the tau deposition. (**D**) Higher-order tau increased from Aβ- and Aβ+ subjects, or from unimpaired Aβ- to unimpaired Aβ+ to impaired Aβ+ stages. (**E**) Higher-order tau was significantly correlated with decreased gBOLD-CSF coupling strength in the whole cohort, Aβ+ subjects and impaired Aβ+ subjects, similar to Fig. 1. Each error bar represents the standard error of the mean; each point in **B** and **E** represents one DKT parcel and one subject, respectively; the sample sizes of each subgroup are shown by numbers on the bars of the bar plots hereafter.

In the PG map, the DMN and frontoparietal network (FPN) best capture regions at the high-order end of the hierarchical gradient and thus were extracted as a higher-order mask (**Fig. 4C**). Tau deposition in these higher-order regions was greater in Aβ+ compared to Aβ- subjects. There was also greater tau burden in impaired Aβ+ individuals compared to unimpaired Aβ+ individuals, who in turn showed greater tau burden compared to unimpaired Aβ- individuals (all *p <* 0.05, two-sample t-test; **Fig. 4D**), suggesting that tau in higher-order regions tracks with AD severity. Consistent with **Fig. 1**, reduced gBOLD-CSF coupling was associated with higher-order tau across the whole cohort, Aβ+, and impaired Aβ+ groups (both *r >* 0.23, *p <* 0.030; **Fig. 4E**).

### High-order tau deposition may be attributed to coupling-related brain propagations dynamics

Further analyses focused on identifying the brain dynamics underlying the gBOLD-CSF coupling and investigating the association between these dynamics and tau or cognitive measures. Global brain activation often takes the form of infra-slow (< 0.1 Hz) propagating waves/events between lower- and higher-order cortical regions (very close to the higher-order tau mask noted above), along the PG direction (*47, 48, 57*). We thus extracted the dynamic propagating waves of cortical activation around several specific gBOLD peaks following a previously described method (*47*). The gBOLD propagating waves were manifested as tilted bands in the time-position graphs showing brain activation in the PG-sorted ROIs transiting along the PG direction between lower-order (e.g., sensory-motor [SM] networks) and higher-order (e.g., DMN) regions (**Fig. 5, A** and **B,** first column). The time-position graphs for the lower-order sensory-motor to higher-order association (Low-to-High order or L-H; averaged across N = 621 events from 115 subjects) and the High-to-Low order or H-L (N = 576; event-based mean) propagations, as well as corresponding spatial patterns (within the -7.2-sec to +7.2-sec windows around gBOLD peaks), are shown in **Fig. 5, A** and **B**, respectively. In addition to the topographic information about progression of waves from Low-to-High order cortex and the reverse, the bands demonstrate reduced intensity at higher-order cortex, compared to the lower-order end. For each subject, the occurrence rates (the number of propagating events during the rsfMRI scanning within approximately 10 minutes) of the bidirectional propagation were summed. Subjects with more propagation events (larger occurrence rates) had less tau deposition in the higher-order cortices (*r =* - 0.23, *p =* 0.015; **Fig. 5C**), stronger gBOLD-CSF coupling (*r =* - 0.21, *p =* 0.022; **Fig. 5D**; especially for the Aβ+ and impaired Aβ+ groups shown in **Fig. S9**), and higher MoCA scores (*r =* 0.20, *p =* 0.040; **Fig. 5F**; consistent with the strong association between gBOLD-CSF coupling and MoCA in **Fig. S10**). Neither of these close links between propagation occurrence and tau, coupling, or MoCA was affected by the number of gBOLD peaks over the rsfMRI time-course (**Fig. S11**), suggesting the important role of specific gBOLD peaks with propagating waves in tau pathology. We found no significant correlation between the propagation frequency and thickness in higher-order association regions (*r* = 0.10, *p* = 0.27), suggesting a weak link between the brain structure and the dynamic propagating activity.

**Fig. 5.**
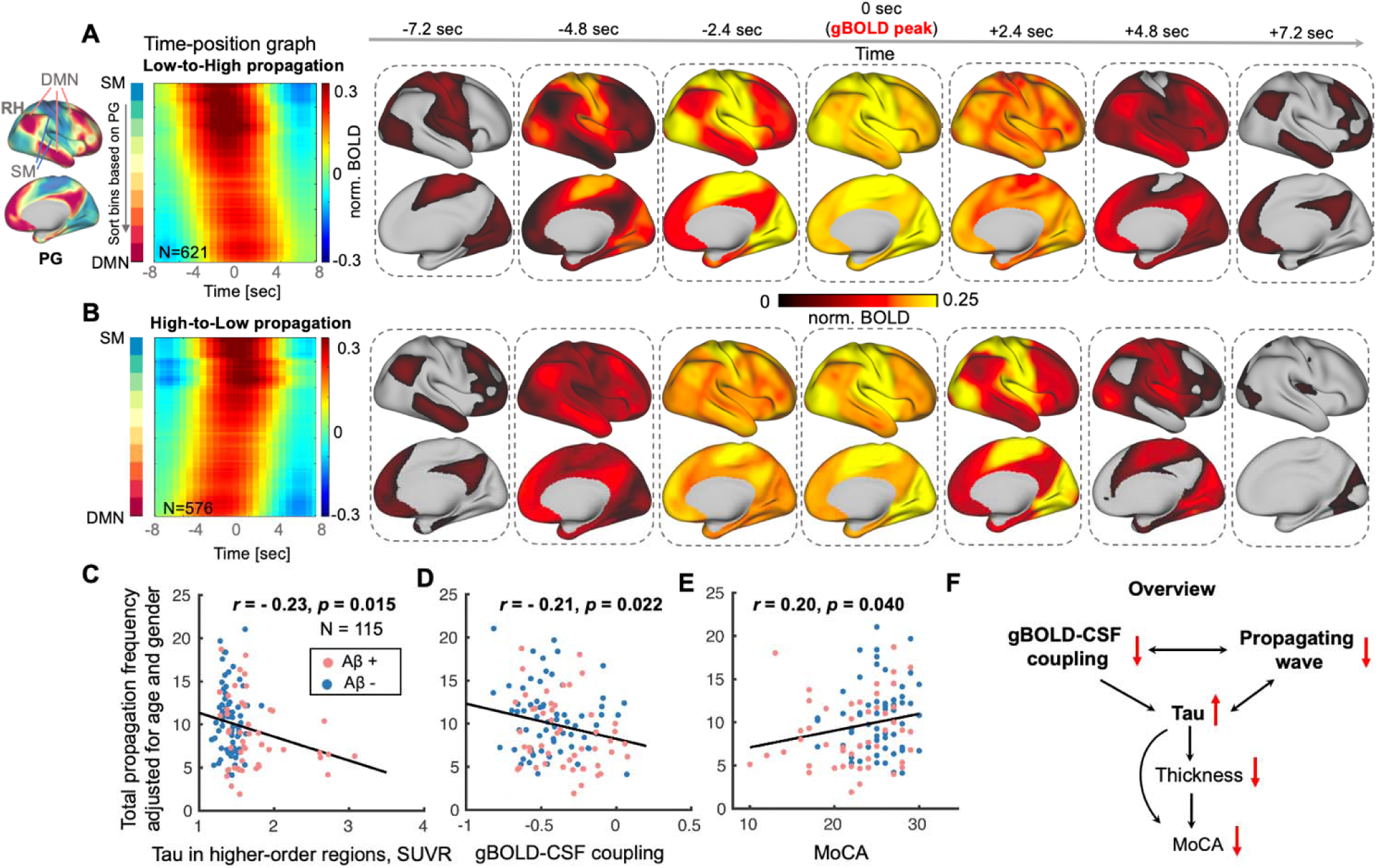
Brain propagation dynamics involved with higher-order cortices are linked to tau, coupling metrics, and cognitive measures. (**A-B**) Propagations between lower-order sensory-motor (SM) and higher-order association cortex were identified near gBOLD peaks as previously described (*47*). The two group-mean time-position graphs are shown as two types of continuous titled bands reflecting brain activation gradually transiting from low-to-high (L-H) order or in the opposite direction, which are shown as spatial pattern changes in the right seven columns. (**C-E**) The frequency of all identified propagation events (sum of the propagation frequency of both low-to-high and high-to-low (H-L); extracted within an equal length of fMRI time-series for each subject; adjusted for age and sex) decreased with the increase of tau deposition in higher-order regions, weaker gBOLD-CSF coupling, and the MoCA decline, across all subjects (all *p* < 0.040). This propagation-coupling relationship was also strong for Aβ+ and impaired Aβ+ subjects (**Fig. S9**; both *r* < - 0.26, both *p* < 0.068), similar to Fig. 1 and 2. (**F**) An overview summarizes the links among coupling metrics, dynamic propagation, and AD markers including thickness, tau, and MoCA. It shows that the glymphatic function-related gBOLD-CSF coupling and dynamic propagating waves modulate tau deposition, cortical thickness, and the cognitive changes, while the propagation frequency and thickness (also in higher-order association regions) was not significantly correlated (*r* = 0.10, *p* = 0.27). Each point in scatter plots represents one subject.

We summarized the multiple associations among gBOLD-CSF coupling, propagation, tau, thickness, and MoCA in **Fig. 5F**. As depicted in this schematic, global brain activity-related coupling metrics, indicative of glymphatic function, and associated dynamic propagating waves modulate cortical tau, thickness, and ultimately affect cognitive performance.

### Relationship between coupling weakening and reduced one-way propagations depends on AD stages

We finally explored the changes of cortical propagations in each of two directions across AD stages and whether the propagation of cortical activity would account for the strength of gBOLD-CSF coupling. The summed occurrence rates of both bidirectional propagations and the L-H ones decreased from Aβ- to Aβ+ stages, and from unimpaired Aβ- to unimpaired Aβ+ stages (all *p <* 0.057, two-sample t-test), but not from unimpaired Aβ+ to impaired Aβ+ stages (**Fig. S12**, **A** and **B**). A similar but weaker trend was also found in the direction of H-L propagation.

For the L-H propagation direction, gBOLD-CSF coupling decreased with reduced propagation occurrence across Aβ- and unimpaired Aβ- subjects (both *r <* - 0.30, *p <* 0.016; **Fig. S12C**), while the association was marginally significant across the whole cohort (*r =* - 0.17, *p =* 0.071). In other words, in Aβ- subjects, the coupling-propagation relationship in **Fig. 5D** may be attributed to the reduced L-H propagation. For the H-L propagation direction, the coupling was strongly associated with decreased propagation frequencies only across Aβ+ and impaired Aβ+ individuals (both *r <* - 0.36, *p <* 0.018; **Fig. S12D** [*r =* - 0.30, *p =* 0.040 and *r =* - 0.34, *p =* 0.083 after excluding the outlier sample with most propagation events in the Aβ+ and impaired Aβ+ group, respectively]), which may be related to the decreased high-order DMN activity in Aβ+ stage (*59–62*) hindering the propagation initiated from there.

## Discussion

We show a close association between resting-state global brain activity and tau pathology, including the spatiotemporal pattern of tau deposition, brain atrophy, and cognitive decline. The coupling between global brain activity, quantified by gBOLD, and CSF movement was attenuated and correlated with tau deposition in widespread neocortical regions characterized by relatively later-involved Braak stage ROIs (V-VI and III-IV). This finding was evident in the whole cohort but more striking for the Aβ+ and impaired Aβ+ subjects, as would be expected since tau burden is likely to be greater for these groups in later-involved neocortical regions. Declining glymphatic function, reflected by weaker gBOLD-CSF coupling, was also strongly associated with reduced cortical thickness in the same regions, likely related to the mediating role of tau. With the significant aggregation of Aβ, tau deposition is seen earlier in higher-order cortices over lower-order ones (*15, 16, 55, 56*). This preferential deposition of tau is related to impaired gBOLD-relevant glymphatic function and to the dynamics of propagating waves of brain activation between higher- and lower-order cortical regions, occurring at gBOLD peaks, that resembles the spatiotemporal pattern of tau spreading and explains its inter-subject variability. Together, these results suggest that resting-state global brain activity modulates the stereotyped pattern of tau deposition in neocortex among individuals with elevated Aβ, presumably via its effect on glymphatic clearance.

Pathological tau aggregation has received increasing attention because of its strong relation to brain atrophy and cognitive impairment (*8, 11*). While a majority of recent studies have attributed the stereotyped pattern of tau accumulation over cortical regions to neural activity and anatomical and functional connectivity (*17–26*), other data has repeatedly demonstrated that the brain clearance system affects tau pathology (*27*) through the glymphatic pathway (*30, 31*). The glymphatic pathway clears brain wastes through CSF movement pushing the exchange between CSF and ISF and its solutes, including Aβ and tau (*30, 31*).

Several lines of evidence support the link between resting-state global brain activity and glymphatic clearance. First, global brain activity, measured by gBOLD fMRI signal and whole brain electrophysiology signals (*44, 45, 63*), are coupled with CSF movement (*37–41*), a key determinant of glymphatic flow (*27, 29, 30*). This coupling is particularly striking during sleep (*37*), when glymphatic function can be 20-fold stronger than wakefulness (*29*). More recently, the coupling between gBOLD and CSF inflow rsfMRI signals has been found to be correlated with various AD risk factors, cortical Aβ deposition (*38, 39*), older age (*41*), and cognitive decline in AD and Parkinson’s disease (PD) (*38, 40*), further supporting the relationship between global brain activity and glymphatic function. Second, neuronal firing cascades that underlie global spontaneous brain events and gBOLD signal are often accompanied by modulation of sympathetic outflow, including cardiac and respiratory pulsations (*64–68*), heart rate variability (*69*), and pupil size (*70–72*). The sympathetic activity could not only facilitate peri-arterial CSF movements via arterial constriction (*66, 73*) but also arouse slow (< 0.1Hz) modulations of cardiac and respiratory pulsations, considered as the major driving forces of glymphatic CSF movement (*74–76*). Third, the intrinsic subcortical vasoactive pathways (*73*), particularly the basalo-cortical projections (*77*) relevant to the cholinergic system and astrocytes, are involved with modulation of global brain activity on vascular tone (*66*). Importantly, while a small proportion of perivascular neurons have direct contact with the vessel wall, most abut astrocytic endfeet (*73*) constituting the astroglial aquaporin-4 (AQP4) channels that facilitate glymphatic flow (*30, 78*). A recent study suggested that intrinsic large astrocytic Ca^2+^ spikes were coupled with the negative gBOLD peaks (*79*). In short, global brain activity and specific neural and physiological factors play a critical role in supporting glymphatic flow and thus affect the AD progresses via moderating the “accumulation-removal balance” of toxic proteins, such as Aβ and tau.

Recent studies in animal models have repeatedly demonstrated the role of glymphatic function in tau aggregation (*31, 80–82*). Using intracortical injection of human tau into mice, a previous study tracked the tau clearance pathway and found that tau can be removed by CSF flow via glymphatic routes, especially for those with traumatic brain injury (TBI), a risk factor for tau aggregation (*80*). Further investigations on tau and glymphatic function suggested that tau is cleared from brain by an AQP4- dependent mechanism (*81, 82*). For example, a recent mouse study has suggested that the impaired CSF-ISF exchange and AQP4 polarization, especially using an AQP4 inhibitor, in the glymphatic system could exacerbate or even induce pathogenic accumulation of tau (*81*). The same study further showed an inverse association between glymphatic function and tau deposition in the healthy mouse cortex (*81*). In addition, the glymphatic system was hypothesized to affect the cell-to-cell propagation of tau in brain (*83*), since tau can be secreted and taken up by both neurons and glia (*84, 85*), and tau secretion to the extracellular space plays an important role in intracellular tau spreading (*80, 86, 87*). Beyond these mouse studies, a recent human study identified the relationship between reduced whole-brain glymphatic activity, assessed with diffusion tensor image analysis along the perivascular space (DTI-ALPS), and the deposition of tau using PET along with cognitive decline (*88*). All these studies support our findings showing the strength of gBOLD-CSF coupling reflecting glymphatic function decreased with tau deposition and cognitive impairment.

It is not surprising that tau mediates the close association between glymphatic function and cortical thickness. Our results showing a close association between coupling and atrophy is consistent with a recent study showing that the impaired glymphatic function in TDP-43 transgenic mice, mimicking the pathology of amyotrophic lateral sclerosis (ALS), was accompanied by neocortical atrophy (*89*). More importantly, middle-aged AQP4 knock-out mice showed elevated tau in both CSF and hippocampus, as well as severe brain atrophy with thinner cortices and hippocampus (*82*). This brain atrophy was attributed to neuronal loss, including the reduction of dentate granule cells and pyramidal cell layer neurons in the piriform cortex, presumably modulated by tau aggregation induced by the AQP4 deficiency (*82*).

AD-related tau predominantly deposits in higher-order cognitive networks over lower-order sensory-motor networks (*15*), consistent with our finding that tau deposition follows this brain hierarchy. This is compatible with significant evidence for early tau accumulation regions resembling connectivity-defined networks (*19–23*). Moreover, global brain activity, profoundly affecting glymphatic function, shows a sensory-dominant pattern opposite to early tau deposition (*45*), and propagates as waves traveling between higher- and lower-order cortical regions (*47*). Interestingly, the identified propagating waves in our study displayed higher intensity in lower-order SM cortex, in both directions whether “Low-to-High” and “High-to-Low” (dark red elements in the time-position graphs, **Fig.5 A** and **B**), consistent with the hypothesis that the stronger brain activity during propagating waves in the lower-order regions may prevent tau deposition there. Consistent with this notion, a previous study suggested that ADNI participants in the earliest stages of AD with stronger SM activity during propagation appeared to have higher CSF Aβ42 (**Fig.5** in (*39*)). These notions thus not only shore up the tight link between the glymphatic function, reflected by the gBOLD-CSF coupling, and the propagating waves at gBOLD peaks, but also support the critical role of dynamic factors in the spatiotemporal pattern of tau deposition. It is worth noting that increased neuronal activity stimulates the release of tau and enhances tau accumulation (*18*). However, it has not been previously shown that dynamic brain activity, i.e., propagating waves, has an impact on the propagation of tau pathology. The findings presented here provide initial evidence that the frequency of propagation waves is correlated with tau burden in higher-order brain regions. In addition, recent dynamic functional connectivity analyses have repeatedly shown that higher order association regions, including the DMN and FPN, play prominent roles in AD progression (*90, 91*), especially the strong correlation between elevated tau accumulation and the declined dynamic activity in DMN and its posterior regions (*91*), which further confirms the effect of the brain dynamics in high-order DMN on AD pathology. Our findings imply that elevated tau deposition in higher order cortex may be related to failure of initiation of H-L propagating waves or failure of L-H propagating waves to reach association cortex, and the weaker activation there compared to the lower-order SM network. Our data warrants future follow-up studies to validate and extend the findings. Together, the dynamic global brain activity may affect the spatiotemporal pattern of tau deposition.

Exploring dynamic brain activity, particularly the directional propagating waves that drive glymphatic function, is of high scientific interest, but complicated by the fact that glymphatic flow and brain activity are difficult to record simultaneously *in-vivo* in the human brain. Interestingly, the present study not only linked the occurrence of propagation waves to gBOLD-CSF coupling, reflecting glymphatic strength across the entire cohort of subjects, but further identified that propagation in two opposite directions could account for the glymphatic function in distinct populations. Specifically, among (unimpaired) Aβ- subjects, coupling reductions can be explained by the occurrence of L- H propagation decrease, which is also related to significant increase of cortical Aβ deposition (i.e., from Aβ- to Aβ+). Previous findings from the ADNI cohort have also demonstrated that the reduction of CSF Aβ-42 is accompanied by both reduced gBOLD-CSF coupling and weaker L-H propagation among Aβ- subjects with significantly lower CSF Aβ-42 (*39*). On the other hand, H-L propagation may account for coupling changes in later AD progression (i.e., across impaired Aβ+ subjects; **Fig. S12D**), when glymphatic function reflected by the coupling metrics was closely associated with tau deposition, suggesting the potential link between H-L propagation and tau pathology (although the frequency of H-L propagation may not be sensitive enough to account for the tau in high-order regions; *r =* - 0.14, *p =* 0.15). This might result from the decreased DMN activity often occurring at the Aβ+ stage (*59–62*) and interfering with the start of H-L propagating waves there. However, these interpretations should be further confirmed with a more precise quantification of dynamic brain activation.

Our current study systematically demonstrates that gBOLD-CSF coupling reflecting glymphatic function and underlying propagation dynamics is strongly related to tau deposition and further affects cortical thickness and cognitive function. Previous evidence has repeatedly reported that tau deposition reduces cortical thickness (*12, 92*) and further leads to cognitive decline (*8, 93*). The current study first links the dynamic global brain activity and relevant glymphatic clearance to cortical tau and then suggests the effect of glymphatic function on brain atrophy and cognitive change through the mediating role of tau. It is worth noting that a recent single neuron recording study linking global brain activity to the memory system has shown that a spiking cascade of widespread neurons featuring sequential activation, similar to the global brain dynamics, strongly modulated the occurrence of hippocampal sharp-wave ripples (*70*) that play a critical role in memory consolidation (*94, 95*). This suggests the close link between resting-state global brain activity and cognitive function, consistent with our finding that the MoCA score is related to both gBOLD-CSF coupling and the propagating waves.

There are a few limitations of the present study. First, we used a cross-sectional analysis and evaluated the spatiotemporal pattern of tau spreading through its between-group differences (i.e., Aβ+ vs. Aβ-). A longitudinal design should be used to test tau accumulation rates linked to fMRI-based coupling or propagating waves. Second, the global propagating waves between higher-order association regions and lower-order sensory-motor networks is one type of propagation event at some gBOLD peaks but do not thoroughly account for the dynamic activations underlying global brain activity or its relevant glymphatic movements. Future studies on an independent dataset with a larger sample size are needed to validate and extend our findings on the global brain dynamics underlying glymphatic flow.

In summary, this study provides initial evidence that glymphatic function, reflected by the coupling between global brain activity and the CSF movement signals, is closely associated with tau pathology in people with elevated Aβ deposition. Global dynamic brain activity is related to glymphatic function and may in part explain the preferential exposure of higher-order cognitive brain regions to tau deposition.

## Materials and Methods

### Participants and data

We included 115 participants from the ADNI-3 project according to the availability of tau-PET (18F-Flortaucipir [FTP]; AV-1451), Aβ-PET (either florbetaben [FBB] or florbetapir [FBP]), rsfMRI (TR = 0.607 sec sessions only), and structural MRI (cortical thickness) data. Participants included 4 different disease conditions defined by ADNI (http://adni.loni.usc.edu/study-design/): 6 AD patients, 42 with mild cognitive impairment (MCI), 5 subjects with subjective memory concern (SMC), and 62 healthy controls. We further categorized the participants into “impaired” (AD and MCI) and “unimpaired” (SMC and control) groups based on their clinical diagnosis. Two groupings were applied for our analyses: 1) Aβ+ and Aβ- subjects; 2) unimpaired Aβ-, unimpaired Aβ+, and impaired Aβ+ subjects (20 impaired Aβ- subjects were used to augment the sample size of the Aβ- group in the comparison of fMRI-based glymphatic or propagation measures with Aβ+ group). We also included participant demographics (age and sex) and cognitive assessment, i.e., Montreal Cognitive Assessment (MoCA) scores in our study. All participants provided written informed consent. Ethical approval from the individual institutional review board (IRB; http://adni.loni.usc.edu/wp-content/uploads/2013/09/DOD-ADNI-IRB-Approved-Final-protocol-08072012.pdf) has been granted to the investigators at each ADNI participating site. All the ADNI data were collected per the principles of the Declaration of Helsinki.

Measurement of Aβ-PET, tau-PET, rsfMRI, cortical thickness, and MoCA were obtained from the same study visit (the time interval between pairwise modalities was no more than 183 days [∼ 6 months] (*96*)). The file “UC Berkeley – AV1451 PVC 8mm Res Analysis [ADNI2,3] (version: 2023-02-17)” was used to provide the tau-PET SUVR (*97–100*). Both “UC Berkeley – AV45 8mm Res Analysis [ADNIGO,2,3] (version: 2023-02- 17)” and “UC Berkeley – FBB 8mm Res Analysis [ADNI3] (version: 2023-02-17)” were used to provide the Aβ-PET SUVR (*51, 52, 101*). MoCA score was also directly acquired from ADNI as “Montreal Cognitive Assessment (MoCA) [ADNIGO,2,3]”. All these data, as well as the rsfMRI and cortical thickness, are publicly accessible on the ADNI website (http://adni.loni.usc.edu/).

The use of de-identified data from ADNI has been approved by the University of California Berkeley and also strictly followed the ADNI data use agreements.

### Image acquisition and preprocessing

All rsfMRI was acquired at 3 Tesla MR scanners from multiple ADNI participating sites following a unified protocol (http://adni.loni.usc.edu/methods/documents/mri-protocols/). The MRI data used in the current study was collected in Siemens MRI scanners (Siemens Medical Solutions, Siemens, Erlangen, Germany). Each MR imaging session began with a T1-weighted (T1w) MPRAGE sequence (flip angle = 9°, spatial resolution = 1 × 1 × 1 mm^3^, echo time [TE) = 3.0 ms, repetition time [TR) = 2,300 ms), which was used for cortical thickness, anatomical segmentation, and registration (see details in http://adni.loni.usc.edu/methods/documents/) (*102*). During rsfMRI acquisition, 976 fMRI volumes were collected with an echo-planar image (EPI) sequence with TR/TE=607/32 ms (flip angle = 50°, spatial resolution = 2.5 × 2.5 × 2.5 mm^3^, slice thickness = 2.5 mm; see details at: http://adni.loni.usc.edu/methods/documents/).

PET imaging was acquired according to standardized protocols at each ADNI site. FTP-PET data were acquired from 75-105 minutes post-injection of 10 mCi tracer. FBP-PET and FBB PET data were acquired from the 50-70 minutes post-injection of 10 mCi tracer and from the 90-110 minutes post-injection of 8.1 mCi tracer (*52, 103, 104*), respectively (https://adni.loni.usc.edu/wp-content/uploads/2012/10/ADNI3-Procedures-Manual_v3.0_20170627.pdf).

We preprocessed structural MRI using FreeSurfer v7.1 (https://surfer.nmr.mgh.harvard.edu/fswiki/DownloadAndInstall5.3) (*105*) to derive FreeSurfer ROIs (DKT-68 parcel (*58*)) in participants’ native space and extract the parcel-based cortical thickness. Following the previous study (*38*), we preprocessed the rsfMRI data using FSL (https://fsl.fmrib.ox.ac.uk/fsl/fslwiki) (*106*) and AFNI (https://afni.nimh.nih.gov/) (*107*) with a modification of excluding rsfMRI sessions with large head-motion (session-mean frame-wise displacement [FD] larger than 0.6 mm or the maximal FD larger than 3 mm) (*108*). The general procedures for rsfMRI preprocessing include motion correction, skull stripping, spatial smoothing (full width at half maximum [FWHM] = 4mm), temporal filtering (bandpass filter, 0.01 to 0.1 Hz), and the co-registration of each fMRI volume to its anatomical MRI and then to the 152-brain Montreal Neurological Institute (MNI-152) space. Referring to the previous studies (*38, 68*), we also excluded the procedure of motion parameter regression to avoid weakening the gBOLD signal, and removed the first 20 and last 20 volumes for each rsfMRI session to reduce the edge effect from the temporal filtering and to ensure a steady magnetization.

We used the tau-PET SUVR data summarized in “UC Berkeley – AV1451 PVC 8mm Res Analysis [ADNI2,3] (version: 2023-02-17)”, and the Aβ-PET SUVR from “UC Berkeley – AV45 8mm Res Analysis [ADNIGO,2,3] (version: 2023-02-17)” (for 42 subjects) and “UC Berkeley – FBB 8mm Res Analysis [ADNI3] (version: 2023-02-17)” (for 73 subjects) (*51, 97–101*). To generate the PET data in DKT-68 parcels, several preprocessing steps were performed, including image averaging, spatial smoothing, and registration to the (structural) MRI space to extract the tau or Aβ intensity in each DKT-68 parcel (*58*). We then normalized parcel-wise tau with the inferior cerebellar reference region to derive the tau SUVR and further applied partial volume correction (PVC) to reduce the influence from low image resolution and limited tissue sampling (*100*). Regarding the Aβ SUVR, we normalized the FBP or FBB intensity in each DKT-68 parcel using the whole cerebellum reference region.

### The extraction of gBOLD and CSF inflow signals

Following the previous study (*38*), we derived the gBOLD signal by averaging the rsfMRI (Z-normalized) time-series across all voxels in the gray-matter region (see a representative example in **Fig. S1B, upper, green;** corresponding to the signal in **Fig. S1A**, **green**). Specifically, we used the Harvard-Oxford cortical and subcortical structural atlases (https://neurovault.org/collections/262/) to define the gray matter mask, which was then transformed to the original fMRI space of each subject referring to previous studies (*37, 38*). The rsfMRI in individual original space went through the above preprocessing procedures (not including nuisance regression; particularly to avoid the CSF signal regression attenuating the CSF inflow signal) (*38*). The preprocessed rsfMRI signal was averaged in individual gray-matter mask to derive the gBOLD signal for each subject.

To derive the CSF inflow signal, the preprocessed fMRI in the original individual space was averaged across the CSF voxels at the bottom slice of fMRI acquisition following previous studies (*37, 38*). All subjects have their individual CSF masks (**Fig. S1B**, **lower**; corresponding signal in **Fig. S1A**, **purple**) with similar voxel numbers (∼14).

### The coupling between the gBOLD signal and the CSF signal

We also calculated the cross-correlation functions between the gBOLD signal and the CSF inflow signal (by assessing Pearson’s correlation) at the lag of +4.856 sec, where the negative peak of the mean cross-correlation located (see arrow in **Fig. S1C**), to evaluate the gBOLD-CSF coupling for each subject as the previous study (*38*).

### Stage classifications

Two grouping methods were applied for the entire group of subjects. First, subjects were divided into Aβ+ and Aβ- based on the whole cortical SUVR (AV45–Aβ > 1.11 SUVR or FBB–Aβ > 1.08 SUVR; the whole cerebellum as reference region). Second, we further incorporated the diagnosis information and staged the subjects into (cognitively) unimpaired Aβ-, unimpaired Aβ+, and impaired Aβ+ groups (unimpaired: SMC and control).

### Correlating the gBOLD–CSF coupling to cortical A***β***, tau, thickness, and MoCA

The gBOLD-CSF coupling was evaluated using their cross-correlation at the lag of +4.856 sec. We first compared the gBOLD-CSF coupling (adjusted for age and sex) across two sets of stages above, including the pairwise comparison (two-sample t-test) and the linear trend evaluation for unimpaired Aβ- to unimpaired Aβ+ to impaired Aβ+ (ordinal regression).

For each of these 5 stages and the entire cohort, we correlated the gBOLD-CSF coupling (age- and sex-adjusted) with the cortical tau SUVR in each DKT-68 parcel and then mapped to the brain surface (using the WorkBench software [version: 1.5.0; https://www.humanconnectome.org/software/workbench-command]) to see the spatial distribution of correlation coefficients. We further tested the coupling-tau correlation in 4 ROIs, including Braak V-VI ROI, Braak III-IV ROI, temporal meta-ROI, and Braak I ROI across subjects within each of these 5 different groups.

The coupling-tau association was retested after regressing out the subject-wise head motion measure, the mean FD, from the gBOLD-CSF coupling, especially in the Braak V-VI ROI, Braak III-IV ROI, and temporal meta-ROI among the whole cohort, Aβ+, or impaired Aβ+ subjects.

Similar to the above analyses on the coupling-tau association, we also correlated the gBOLD-CSF coupling with the cortical thickness in the same sets of ROIs or MoCA scores (also examined the MoCA-thickness correlation) across the same groups of subjects.

### Determining the mediator role of tau in the coupling-thickness and coupling-MoCA associations

We first averaged tau SUVR and cortical thickness across all subjects and compared their distribution patterns by correlating them across DKT-68 parcels. Similarly, the tau difference between Aβ+ and Aβ- groups (reflecting the tau spreading with AD progression) was spatially compared with that difference in thickness. We further evaluated the inter-subject similarity between tau and thickness in each of the above 4 ROIs across subjects within each group of these 5 different stages.

Given that the gBOLD-CSF coupling may reveal glymphatic clearance for Aβ and tau, as well as the role of tau in predicting brain atrophy (*12, 92*), we then examined the hypothesis that tau mediates the association between gBOLD-CSF coupling and cortical thickness using a mediation analysis (*109*) in the Braak V-VI ROI, Braak III-IV ROI, or temporal meta-ROI among the groups of the whole cohort, Aβ+, and impaired Aβ+, respectively, among which the coupling-tau and coupling-thickness associations were significant in **Figs. 1** and **2**. The possibility that coupling mediated the relationship between tau and thickness was also tested in these ROIs.

Same mediation analysis was further applied to test if the coupling-MoCA association would be mediated by the tau, since tau pathology may lead to cognitive impairment (*8, 110*).

### Relating the mean tau pattern and its cross-stage changes to brain hierarchy

Due to the earlier accumulation in higher-order brain regions than lower-order sensory ones (*15, 16, 55, 56*), we tested whether tau and its cross-stage change followed the brain hierarchy. We thus applied the PG map obtained in (*57*) to quantify the cortical hierarchy and correlated the tau and PG values across cortical parcels. We also assessed such spatial correspondence between PG and tau changes from Aβ- to Aβ+ groups.

### Quantifying tau in higher-order brain regions

We further defined a higher-order cortical mask following a previous study (*39*) to better quantify the tau in higher-order regions, which may reflect the spreading pattern of tau during AD progression. We employed the DMN and FPN defined from a previous study (*111*) as the higher-order association regions and then identified the DKT-68 parcels belonging to the two networks due to the (DKT-68) parcellation of tau.

Averaged tau in higher-order brain parcels was then compared between Aβ- to Aβ+ groups, as well as between the groups of unimpaired Aβ-, unimpaired Aβ+, and impaired Aβ+. We also replicated the analyses in **Fig. 1** and examined the correlation between the gBOLD-CSF coupling and tau in higher-order parcels across subjects in the whole cohort, Aβ+ stage, and impaired Aβ+ stage, in which the coupling-tau relationships in **Fig. 1** were significant.

### Quantifying gBOLD propagating waves and linking it to tau pathology

We followed previous studies (*39, 47*) to project the rsfMRI signal onto the direction of PG (indicative of cortical hierarchy) and derived the time-position graphs over the whole time-series for each subject. We sorted all the cortical voxels based on their PG values and divided them into 70 position bins of equal size. The rsfMRI time-series was then averaged within each bin and thus generated as a time-position graph with the horizontal direction encoding time and the vertical direction representing 70 bins along the PG direction. The time-position graph was cut into multiple segments according to the troughs of the gBOLD signal (*47*). Within the time-position graph for each subject, we identified several continuous and tilted bands representing brain activation transiting from lower-order sensory regions to higher-order regions along the PG direction, as the L-H propagation events; we also detected and extracted the bands showing brain activation transiting along the opposite direction as the H-L propagation events. We followed the previous studies in event detection (see more details in (*39, 47*)). The general procedures included, within each segment of time-position graph, deriving the timing relative to the global (spatial) mean peak (i.e., gBOLD peak), and correlating the relative timings with the position of the corresponding bin (across 70 bins) along the PG direction. We used a correlation threshold of 0.385 (corresponding to the *p*-value of 0.001) to extract the L-H propagations and -0.385 to detect the H-L ones, applied a bin number threshold of 50 to identify these “continuous” bands (i.e., excluding the segments/events with less than 50 local peaks [each bin row could have a “local peak” or not]), and used a (normalized BOLD) threshold of 0.1 to exclude these bin-rows with too weak activation (i.e., regarded as no “local peak”).

The time-position graphs for identified L-H propagation events were averaged (time window from -8-sec to 8-sec; aligned based on the timing of gBOLD peak [0 sec]) across all instances from all subjects to derive the mean time-position pattern of the propagation at this direction. This mean time-position graph was then traced back and projected onto the brain surface to generate the spatial patterns of such propagation. The same approach was applied to derive the mean pattern of the H-L propagation and corresponding spatial patterns.

We also counted the number of identified L-H propagation and H-L propagation events within the full time-position graph (of equal length in the horizontal direction; ∼ 10mins) for each subject as the propagation occurrence rate or frequency for the respective direction. The frequencies of the two distinct directions were further summed to derive the total propagation frequency for each subject.

To investigate the role of dynamic activation propagation in tau, gBOLD-CSF coupling reflecting glymphatic function, cortical thickness, and the cognitive level, we correlated the total propagation frequency (age- and sex-adjusted) with the tau deposition and thickness in higher-order association regions, the gBOLD-CSF coupling, and MoCA score across the whole cohort. We also further examined the propagation-coupling association for each of the 5 different (participants) sub-groups.

To demonstrate the role of Aβ or AD pathology in the frequency of all propagation, the summed frequencies of two directions were also compared between Aβ- and Aβ+ groups, or between unimpaired Aβ-, unimpaired Aβ+, and impaired Aβ+ groups. The frequencies of L-H propagation and the H-L propagation were also compared between the above groups.

To investigate the contribution/relation of propagation at each direction to gBOLD-CSF coupling, we correlated the propagation frequency of each direction with gBOLD-CSF coupling at all (participants) sub-groups, including whole cohort, Aβ-, Aβ+, unimpaired Aβ-, unimpaired Aβ+, impaired Aβ+ groups.

### Statistical analysis

A two-sample t-test was performed for group comparisons on continuous measures, including age, gBOLD-CSF coupling, tau SUVR in higher-order regions, and propagation frequency. We used Fisher’s exact test (*112*) to compare categorical measures (i.e., sex) between stages of Aβ pathology progression. An ordinal regression test was applied to evaluate the linear trend of gBOLD-CSF coupling changes across the unimpaired Aβ-, unimpaired Aβ+, and impaired Aβ+ groups. Pearson’s correlation was used to quantify the relationship between different variables in **Fig. 5F**. The single-level mediation analysis (*109*) was performed to evaluate the role of tau in the coupling-thickness and the coupling-MoCA links, as well as the role of coupling in the tau-thickness relationship. A p-value no more than 0.05 was considered statistically significant.

To test the effect of head-motion on our major results, we regressed out the mean FD (*113*) from gBOLD-CSF coupling and repeated the analyses on coupling-tau association across the groups of subjects in **Fig. 1**.

To examine if the gBOLD peak number drove the close association between the frequency of propagation and tau, coupling, or MoCA in **Fig. 5, C-E**, we correlated the gBOLD peak number with the above three measures across the whole cohort of subjects. We further regressed out the gBOLD peak number from the propagation frequency and re-tested the correlations between the adjusted frequency and tau, coupling, as well as MoCA across the whole cohort, respectively.

## Author contributions

F.H. and W.J.J. conceptualized and designed the study; F.H., J.Q.L., J.Z., T.W., S.M.L., and W.J.J. curated the data and performed analyses; F.H., X.C.,S.L.B., and T.M.H. and W.J.J. designed and performed assays; F.H. and W.J.J. wrote the original draft, and J.Q.L.; F.H., X.C., J.Z., T.W., S.M.L., S.L.B., T.M.H. and W.J.J. reviewed and edited the manuscript; W.J.J. supervised the project; and all authors read and approved the manuscript.

## Data and materials availability

The multimodal data, including subject characteristics, rsfMRI, structural MRI, Aβ-PET, tau-PET, and MoCA measures are all publicly available at the ADNI website upon the approval of the data use application (http://adni.loni.usc.edu/). The ADNI was launched in 2003 as a public-private partnership, led by Principal Investigator Michael W. Weiner, MD. The primary goal of ADNI has been to test whether serial magnetic resonance imaging (MRI), positron emission tomography (PET), other biological markers, and clinical and neuropsychological assessment can be combined to measure the progression of mild cognitive impairment (MCI) and early Alzheimer’s disease (AD). For up-to-date information, see www.adni-info.org. All the code used in the present study are available from the corresponding author upon request.

