## Supplemental figures for "Global brain activity and its coupling with cerebrospinal fluid flow is related to tau pathology"

[†] Data used in preparation of this article were obtained from the Alzheimer's Disease Neuroimaging Initiative (ADNI) database (adni.loni.usc.edu). As such, the investigators within the ADNI contributed to the design and implementation of ADNI and/or provided data but did not participate in analysis or writing of this report. A complete listing of ADNI investigators can be found at: http://adni.loni.usc.edu/wp-content/uploads/how_to_apply/ADNI_Acknowledgement_List.pdf

**Address all correspondence to:**

Feng Han

250 Earl Warren Hall

Helen Wills Neuroscience Institute

University of California, Berkeley

Berkeley, CA 94720

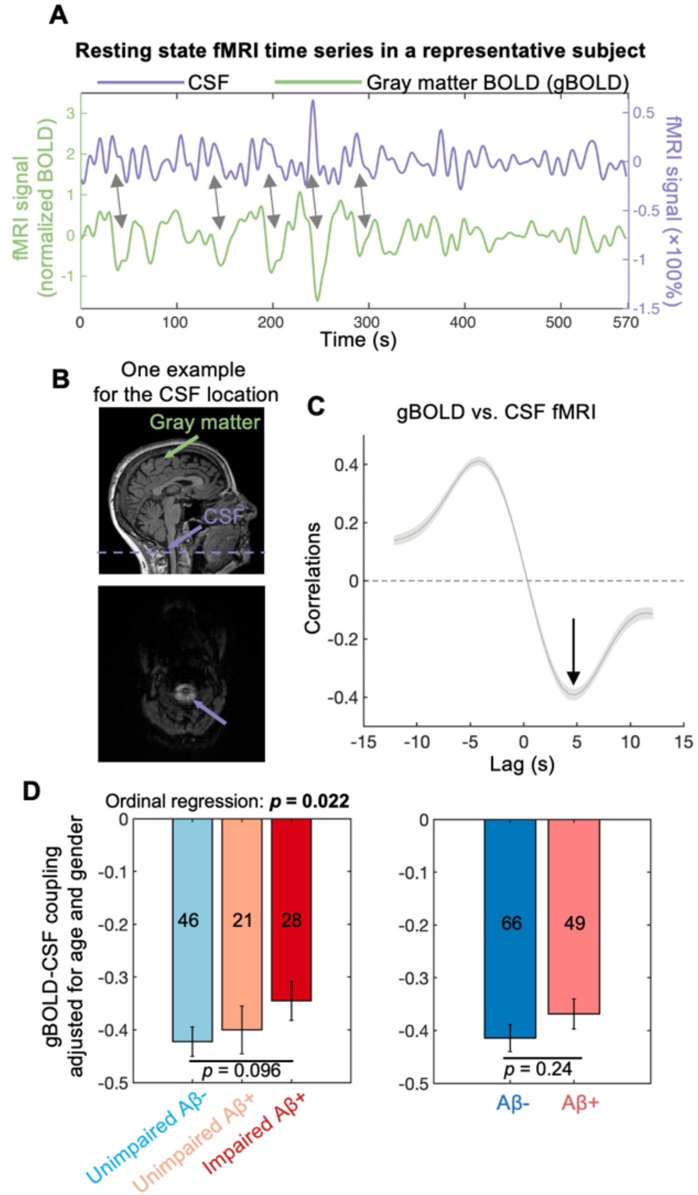


**Fig. S1. gBOLD and CSF inflow signals are coupled.** (**A**) The resting-state gBOLD signal and CSF inflow signal from a representative example showed a negative coupling with a slight time-shift (+4.856 sec; indicating the synchronization with the gray arrows). (**B**) **Upper**: gBOLD signal in (**A**) was extracted with averaging across all gray-matter voxels (green arrow); **Lower**: CSF inflow signal was extracted from the CSF regions at the bottom slice of the fMRI acquisition (purple arrow in the T1w MRI; corresponding to the bottom slice of fMRI acquisition, purple dashed line in the upper panel). (**C**) The averaged cross-correlation functions between gBOLD signal and CSF inflow signal from 115 ADNI-3 subjects. The gray shaded region denoted the one standard error of the mean

(SEM). This mean cross-correlation pattern was similar to previous studies *(37, 38)*. The cross-correlation at the +4.856 sec will be used to measure the gBOLD-CSF coupling for each subject. (**D**) The gBOLD-CSF coupling showed a decrease trend with AD progression (unimpaired Aβ- to unimpaired Aβ+ to impaired Aβ+; ordinal regression, $p = 0.022$), and from Aβ- to Aβ+, although not significant. Each error bar represents the SEM.

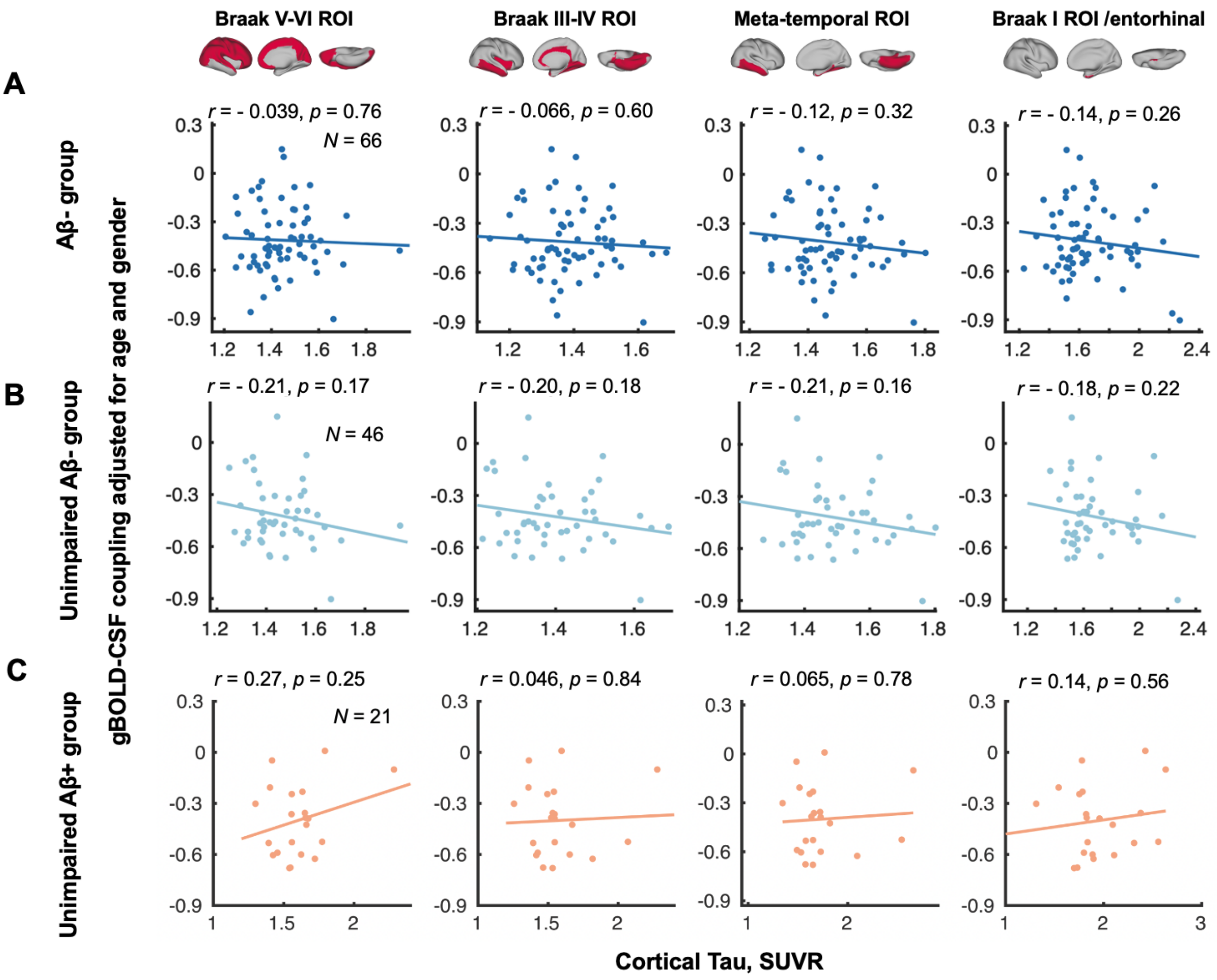


**Fig. S2. gBOLD-CSF coupling is not correlated with the cortical tau across Aβ- or cognitively unimpaired subjects.** The gBOLD-CSF coupling was not correlated with tau in Braak V-VI ROI, Braak III-IV ROI, temporal meta-ROI, or Braak I ROI, across Aβ-, unimpaired Aβ-, or unimpaired Aβ+ subjects.

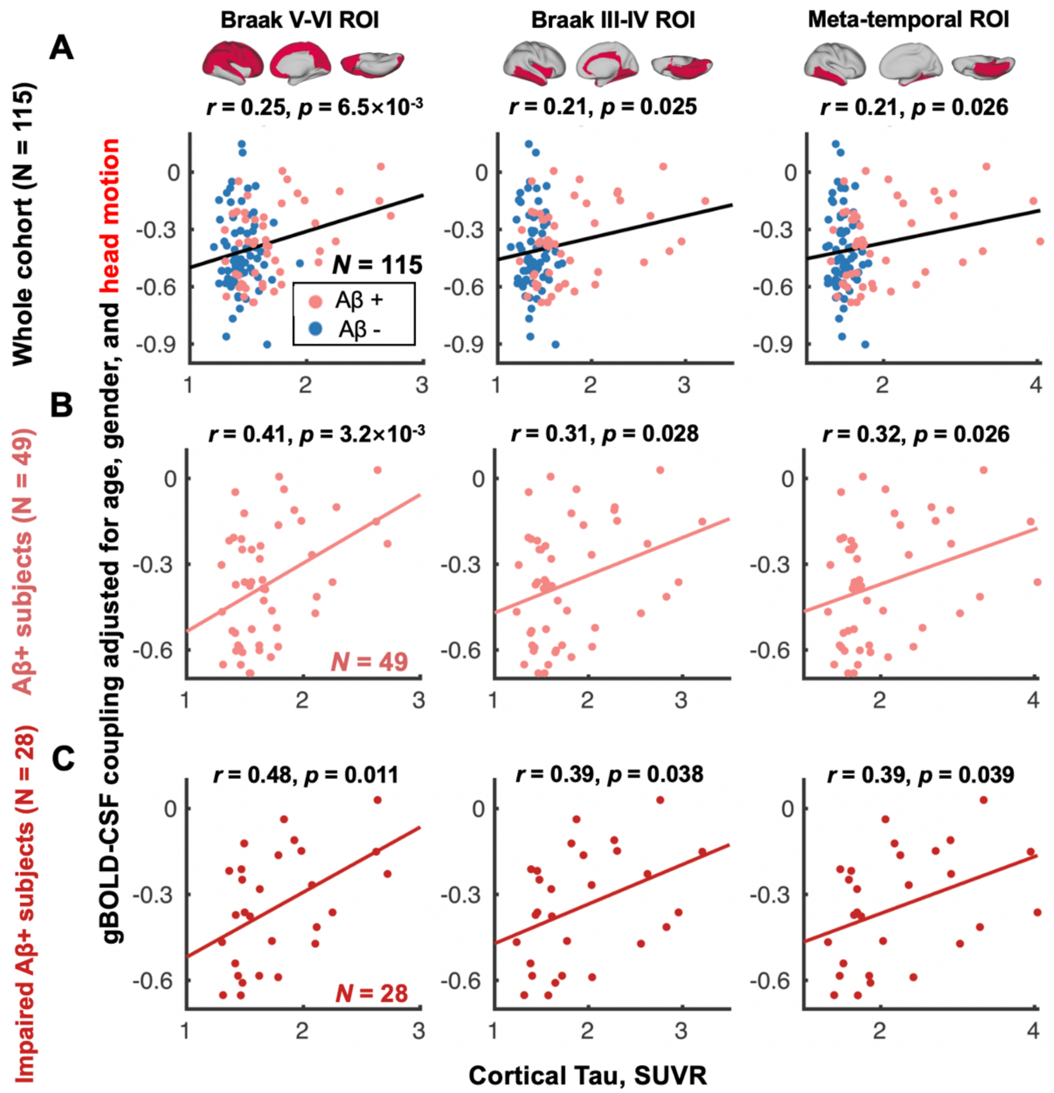


**Fig. S3. The close association between gBOLD-CSF coupling and tau was not affected by head motion.** We repeated the analyses in **Fig. 1**, especially the significant links for Braak V-VI, Braak III-IV, and meta-temporal area, with adjusting the coupling for mean FD, as a subject-wise head motion measure *(68)*. We then found little change before and after the motion adjustment, and these coupling and tau remained strongly correlated after controlling head motion.

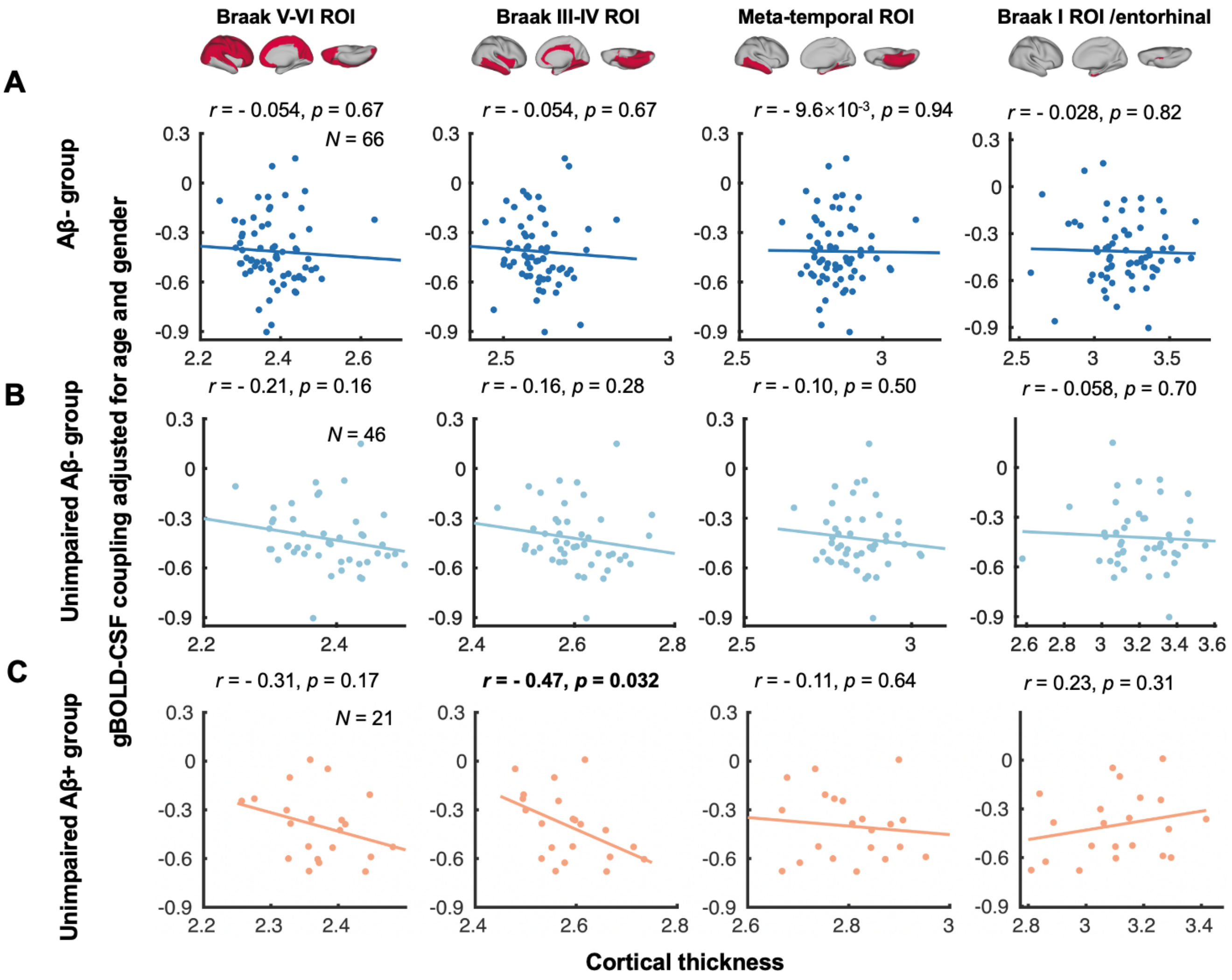


**Fig. S4. gBOLD-CSF coupling is not correlated with the cortical thickness across Aβ- or cognitively unimpaired subjects.** The gBOLD-CSF coupling was not correlated with thickness across Aβ-, unimpaired Aβ-, or unimpaired Aβ+ subjects, in most Braak ROIs or temporal meta-ROI, while the coupling-thickness association was significant in Braak III-IV ROI across unimpaired Aβ+ subjects ($r = -0.47$, $p = 0.032$), which might be related to the rapid tau accumulation in this area among the early Aβ+ population.

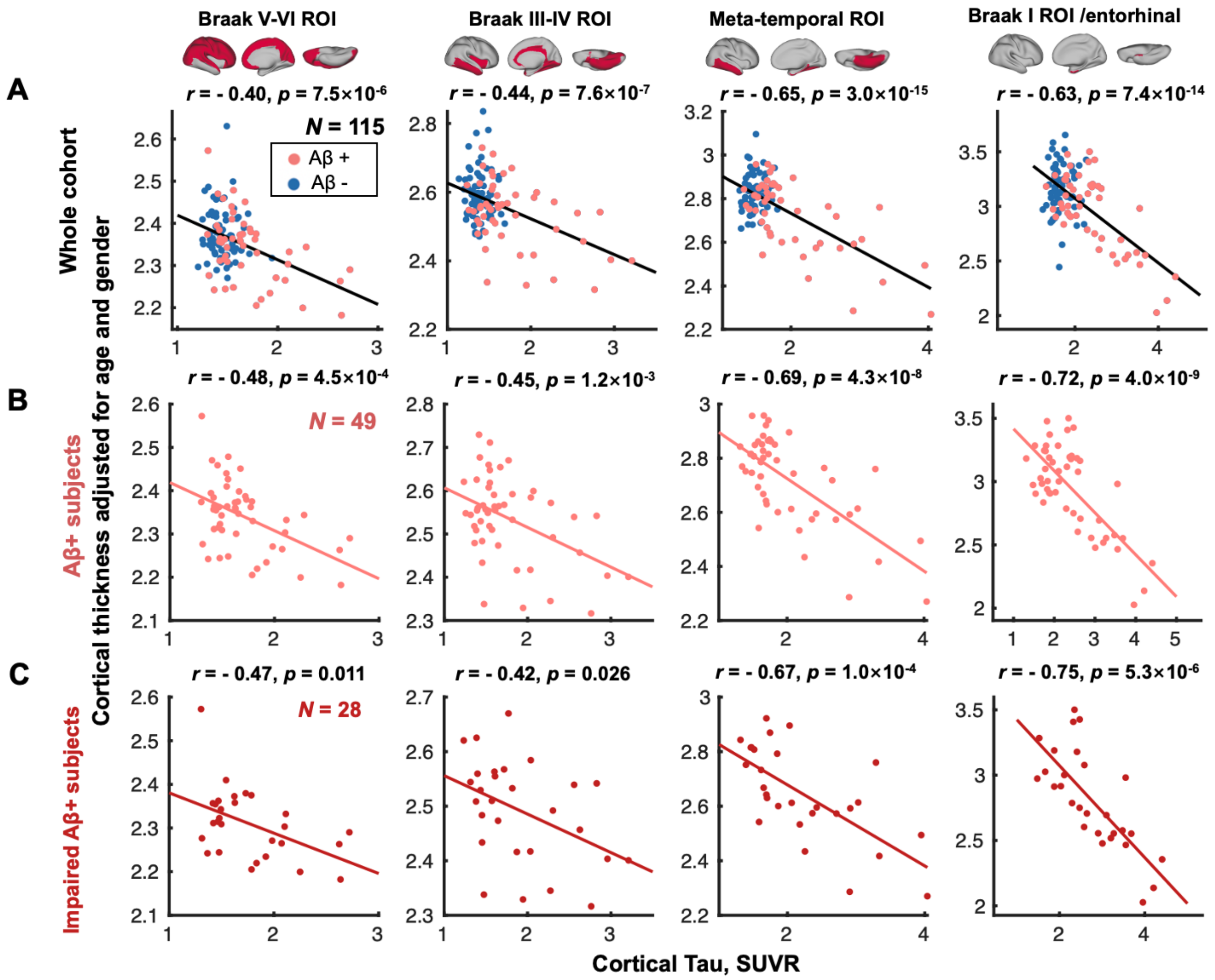


**Fig. S5. Cortical thickness is correlated with the tau in the same regions across the whole cohort and Aβ+ subjects.** (**A**) The cortical thickness was negatively correlated with tau in most cortical regions (i.e., Braak V-VI ROI, Braak III-IV ROI, temporal meta-ROI, and Braak I ROI) across all subjects, i.e., subjects with thinner cortices had more tau deposition (all $r < -0.40$, all $p < 7.5\times10^{-6}$; N = 115). (**B-C**) These associations between cortical thickness and tau were also evident among Aβ+ and/or specifically impaired Aβ+ subjects (all $r < -0.42$, all $p < 0.026$), but not for the rest of the subjects (Aβ-, unimpaired Aβ-, or unimpaired Aβ+ subjects; all $p > 0.13$).

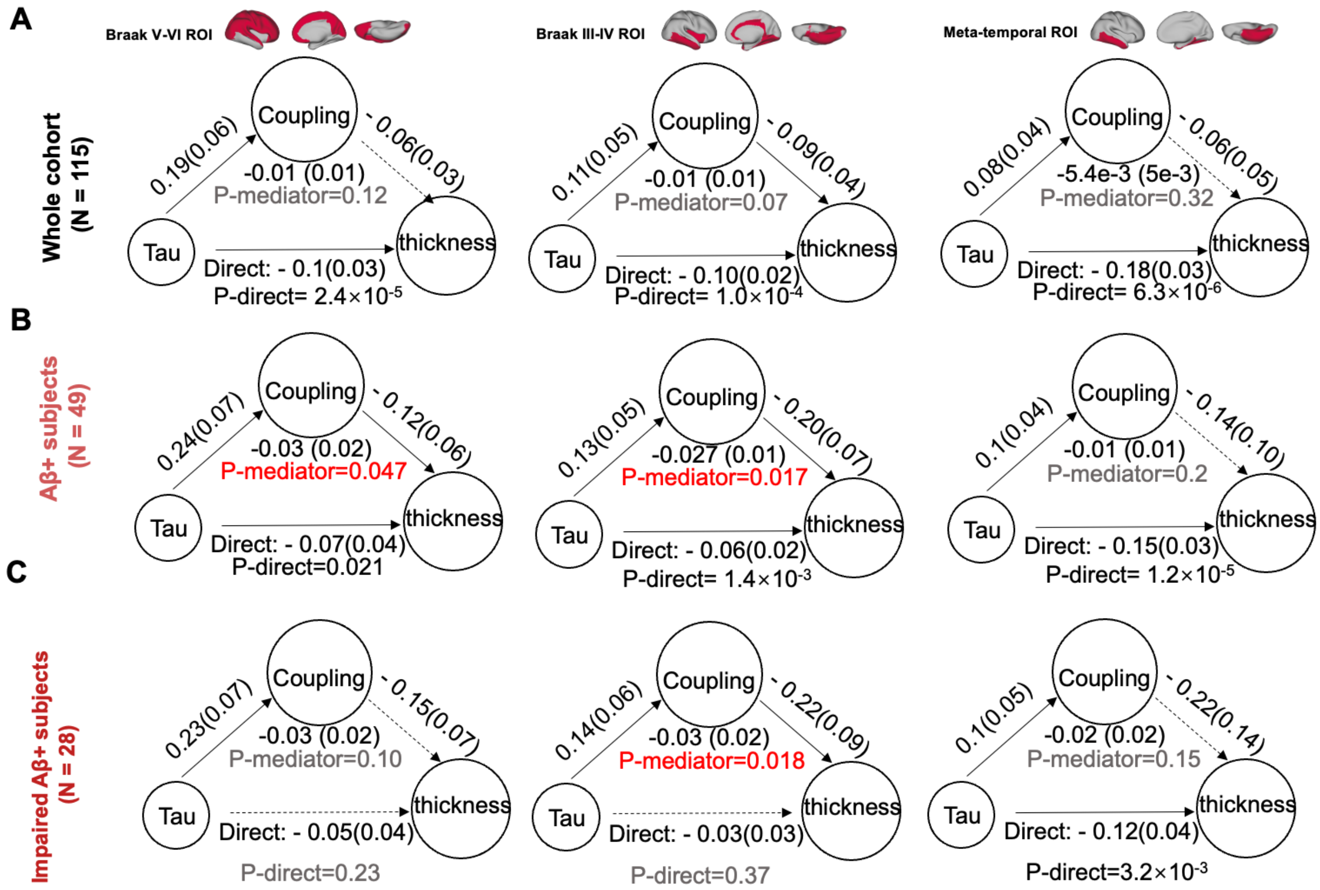


**Fig. S6. The association between tau and thickness may be not strongly mediated by the coupling metrics.** (**A**) Across the entire cohort of subjects, the gBOLD-CSF coupling did not mediate the close association between the tau and thickness in neither cortical region of interest, including Braak V-VI, Braak III-IV, and meta-temporal area. (**B-C**) Although the meditation effects (of coupling) were significant for Aβ+ subjects in the Braak V-VI ROI and Braak III-IV ROI, as well as for impaired Aβ+ ones in the Braak III-IV ROI (all $p < 0.047$), the coupling did not play an evident mediating role among other ROIs or the rest of subjects.

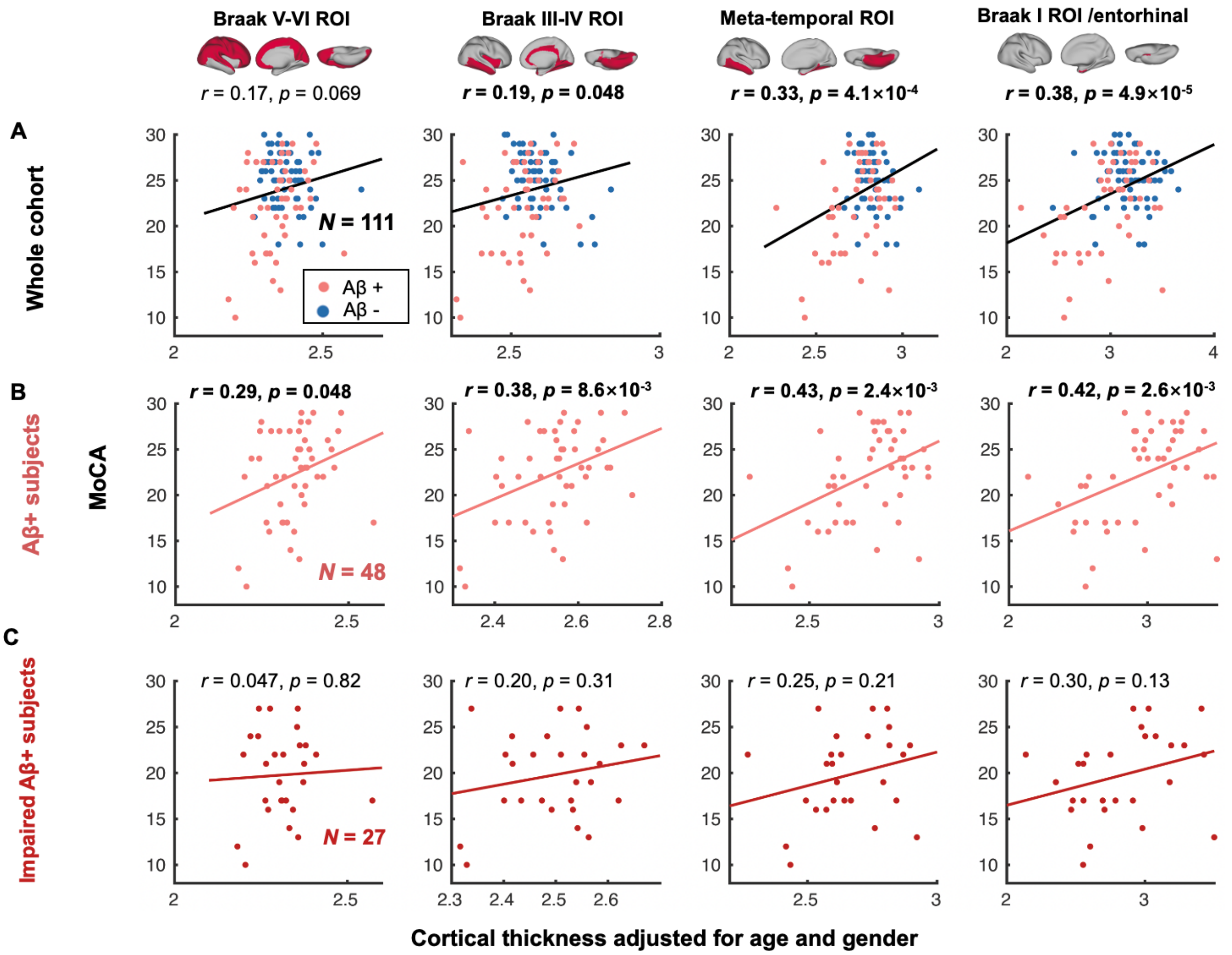


**Fig. S7. MoCA is correlated with cortical thickness in the same regions across the whole cohort and Aβ+ subjects.** (**A**) The MoCA was positive correlated with cortical thickness in most cortical regions (i.e., Braak V-VI ROI [$r$ = 0.17, $p$ = 0.069 marginally significant], Braak III-IV ROI, temporal meta-ROI, and Braak I ROI) across all subjects, i.e., subjects with thinner cortices had worse cognition. (**B-C**) These associations between cortical thickness and MoCA were also evident among Aβ+ subjects (all $r > 0.29$, all $p < 0.048$), while the thickness-MoCA links were not strong in the impaired Aβ+ subjects, presumably relevant to the limited sample size.

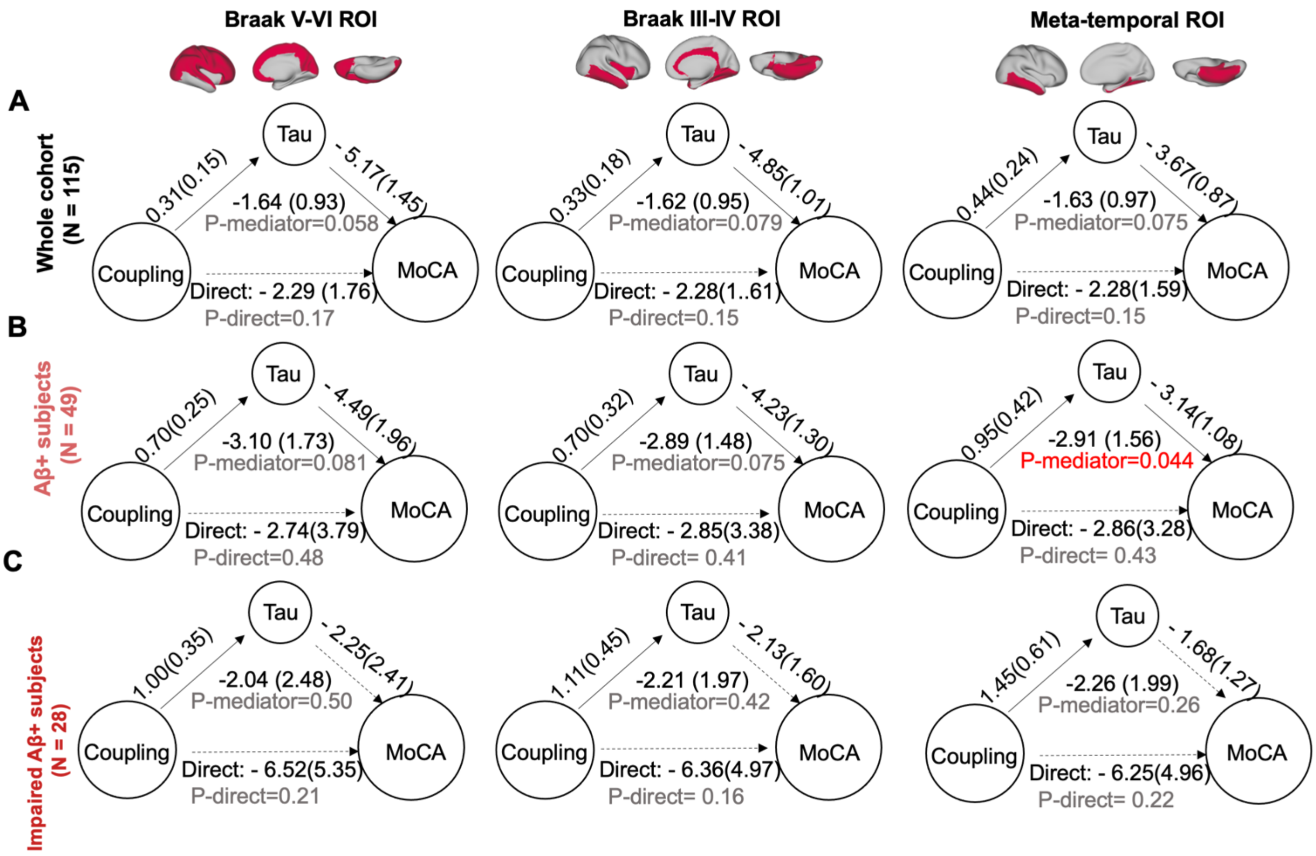


**Fig. S8. The association between coupling and MoCA may be mediated by Tau.** Across the entire cohort of subjects or the Aβ+ ones, the close association between the gBOLD-CSF coupling and MoCA appeared to be marginally mediated by tau in several cortical regions, including the Braak V-VI, Braak III-IV, and meta-temporal area. In particular, the mediation was significant in the meta-temporal regions among the Aβ+ subjects ($p = 0.044$; 3rd column in **B**).

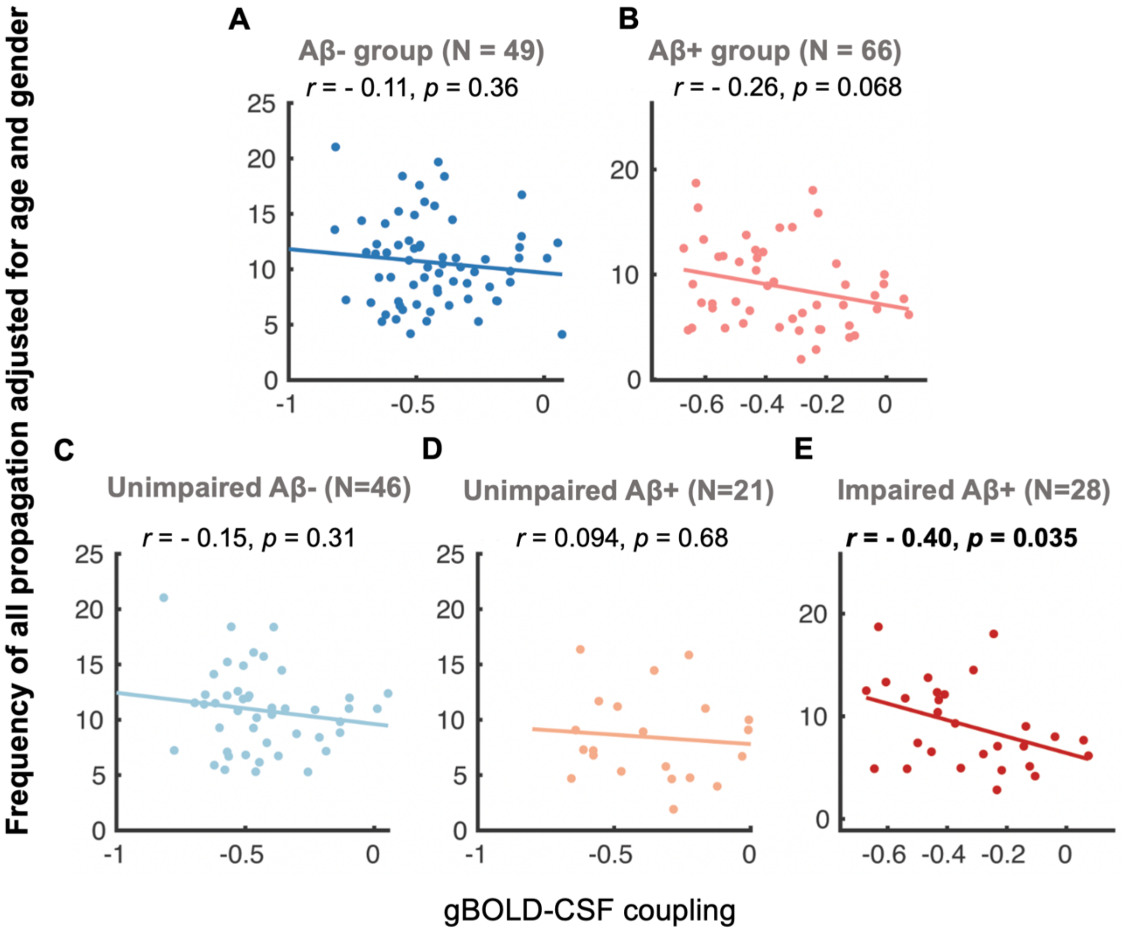


**Fig. S9. The gBOLD-CSF coupling was strongly correlated with summed propagation frequency of two directions across the impaired Aβ+ subjects.** Similar to the strong association between the frequency of all propagation and coupling, the frequency and coupling relationship was strong among Aβ+ (marginally significant, $r = -0.26$, $p = 0.068$) and impaired Aβ+ subjects ($r = -0.40$, $p = 0.035$), not among the rest of subjects. This suggested the propagation between higher- and lower-order regions around the gBOLD peaks may reflect the brain activation dynamics of gBOLD-CSF coupling, at the Aβ+ and impaired Aβ+ stages.

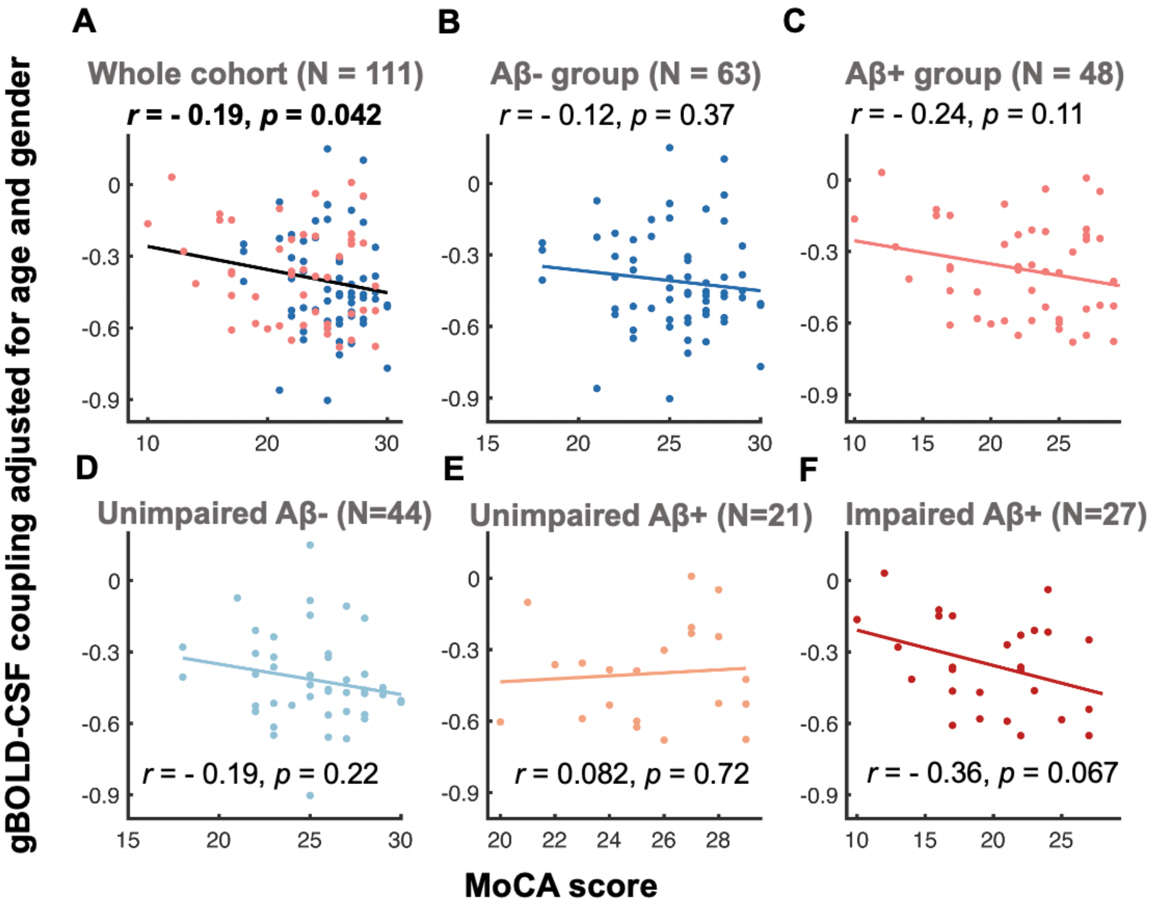


**Fig. S10. The gBOLD-CSF coupling was strongly correlated with MoCA across the whole cohort and impaired Aβ+ subjects.** Across the entire cohort of subjects, subjects with weaker gBOLD-CSF coupling had decreased MoCA scores ($r$ = - 0.19, $p$ = 0.042). This association was also evident in Aβ+ and impaired Aβ+ subjects although not significant (both $p < 0.11$), but weaker among the rest of groups.

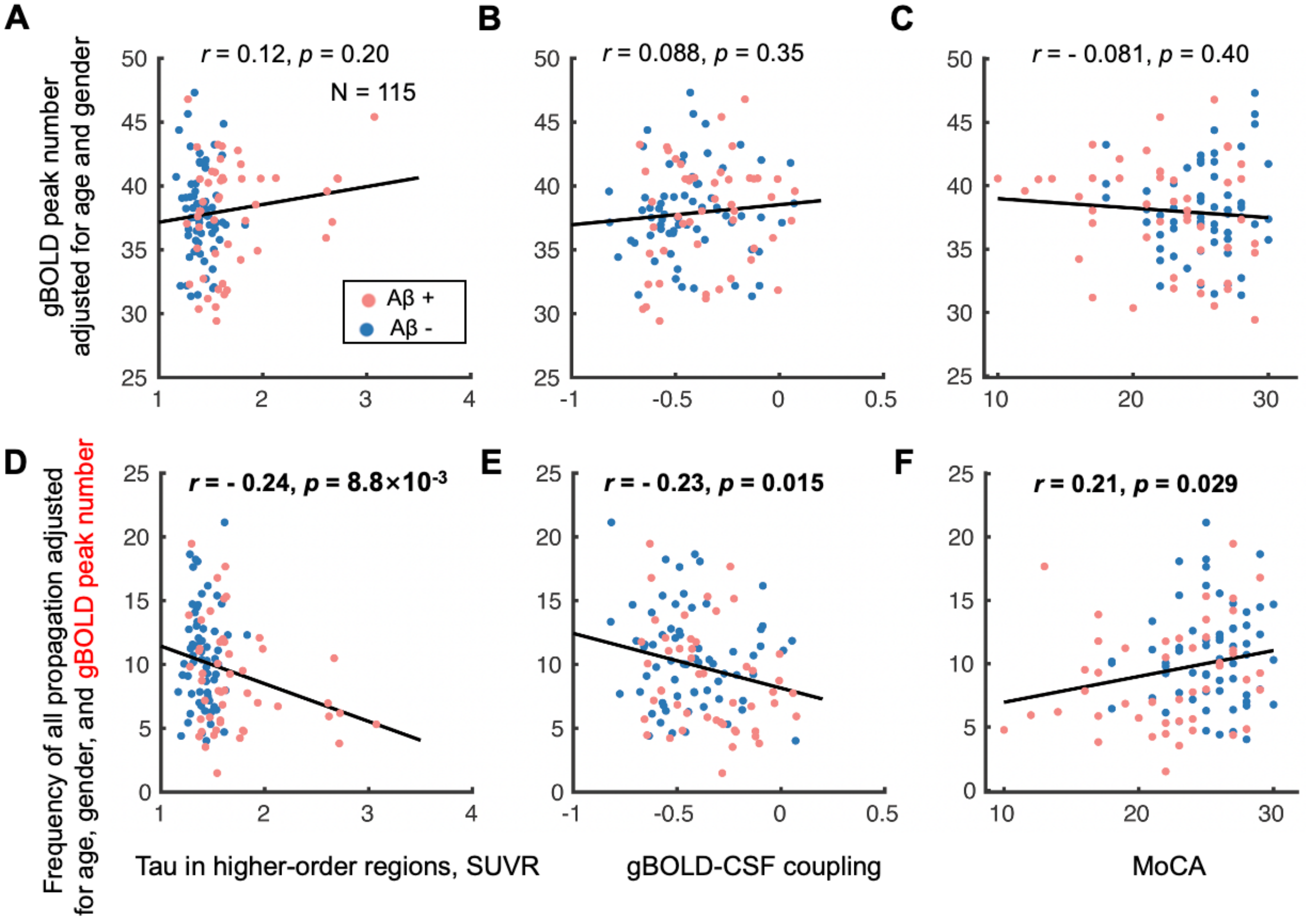


**Fig. S11. The gBOLD peak number has little effects on the links between the number of all propagation events and tau, coupling, or MoCA. (A-C)** The number of gBOLD peaks within the rsfMRI time-series of individual subject was not correlated with the higher-order tau, gBOLD-CSF coupling, and MoCA score. **(D-F)** The association between the total propagation number and tau, coupling, or MoCA remained significant, similar to **Fig, 5, C-E**, after adjusting for gBOLD peak number.

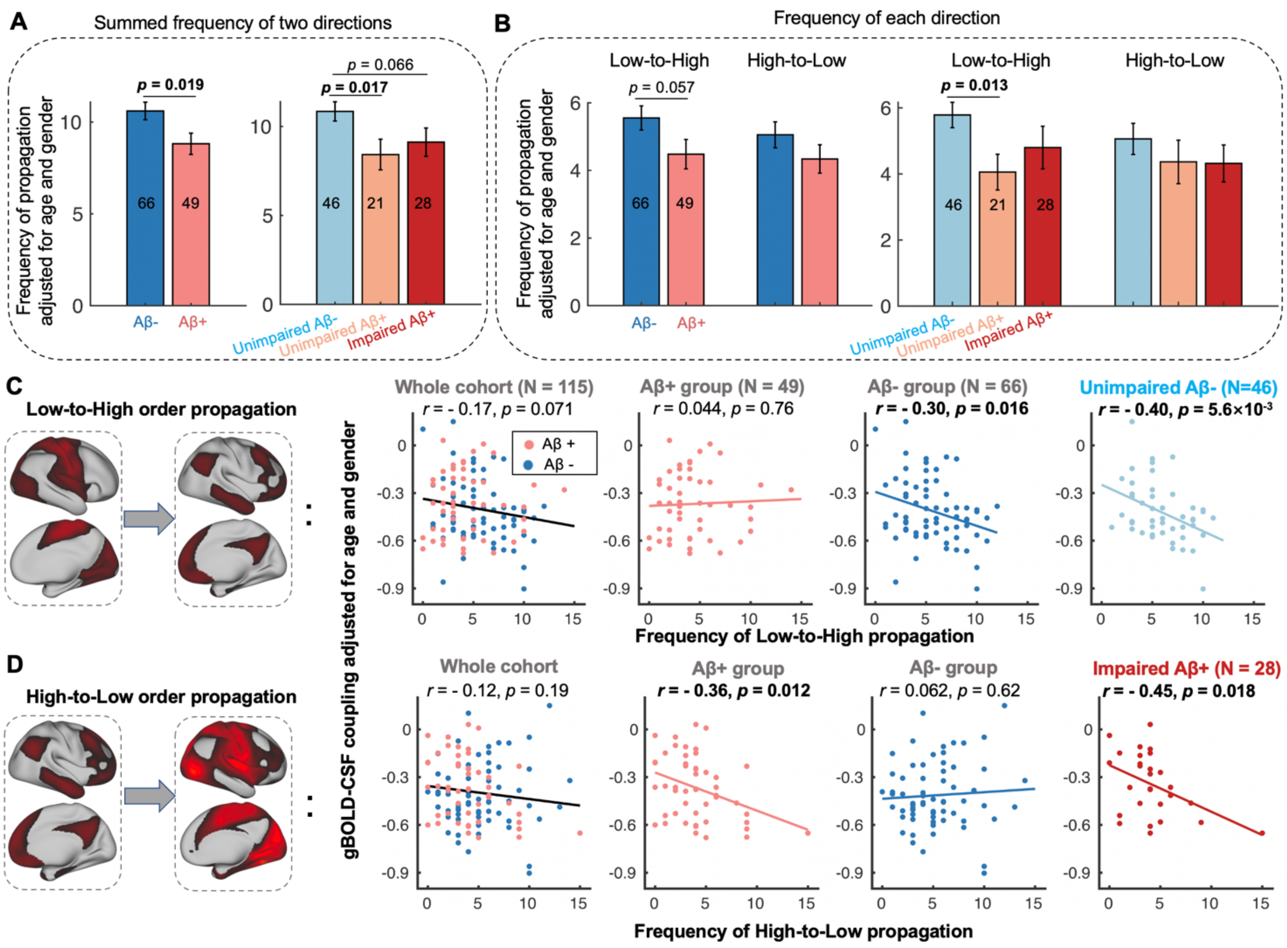


**Fig. S12. Cortical activation propagations of the two different directions and their associations with coupling change with AD progression.** (**A**) Summed propagation frequency for both directions (H-L and L-H) is lower in Aβ+ subjects but not different between unimpaired Aβ+ to impaired Aβ+ stages. (**B**) Group differences in propagation frequency were relatively stronger for the L-H direction, compared to the H-L one. (**C**) For the L-H propagation, the frequency reduced as coupling strength decreased across Aβ- and the unimpaired Aβ- subjects (both $r < -0.30$, $p < 0.016$). (**D**) For the H-L propagation, the frequency reduced as coupling strength decreased across Aβ+ and the impaired Aβ+ subjects (both $r < -0.36$, $p < 0.018$). This frequency-coupling link remained strong after excluding the outlier subject with the most propagation ($r = -0.30$, $p = 0.040$ for the Aβ+ subjects; $r = -0.34$, $p = 0.083$ for the impaired Aβ+ ones). Each error bar represents the standard error of the mean; each point in the scatter plot represents one subject.
